# EXTRA LARGE G-PROTEIN 2 (XLG2) mediates cell death and hyperimmunity via a novel, apoplastic ROS-independent pathway in *Arabidopsis thaliana*

**DOI:** 10.1101/2021.10.08.463358

**Authors:** Elena Petutschnig, Julia Anders, Marnie Stolze, Christopher Meusel, Ronja Hacke, Melina Schwier, Anna-Lena Gippert, Samuel Kroll, Patrick Fasshauer, Marcel Wiermer, Volker Lipka

## Abstract

- Heterotrimeric G-Proteins are signal transduction complexes comprised of three subunits, Gα, Gβ and Gγ, and are involved in many aspects of plant life. The non-canonical Gα subunit XLG2 mediates PAMP-induced ROS generation and immunity downstream of PRRs. A mutant of the chitin receptor component CERK1, *cerk1*-4, maintains normal chitin signalling capacity, but shows excessive cell death upon infection with powdery mildews. We identified *XLG2* mutants as suppressors of the *cerk1*-4 phenotype.
- We generated stably transformed *Arabidopsis* lines expressing Venus-XLG2 and numerous mutated variants. These were analysed by confocal microscopy, Western blotting and pathogen infection. We also crossed *cerk1*-4 with several mutants involved in immunity and analysed their phenotype. Phosphorylation of XLG2 was investigated by quantitative proteomics.
- Mutations in XLG2 complex partners AGB1 and AGG1 have a partial *cerk1*-4 suppressor effect. The *cerk1*-4 phenotype is independent of NADPH oxidase-generated ROS, BAK1 and SOBIR1, but requires PUB2. XLG2 mediates *cerk1*-4 cell death at the cell periphery. Integrity of the XLG2 N-terminal domain, but not its phosphorylation, is essential for correct XLG2 localisation and *cerk*1-4 signalling.
- Our results suggest that XLG2 transduces signals from an unknown cell surface receptor that activates an apoplastic ROS-independent cell death pathway in *Arabidopsis*.

## INTRODUCTION

Heterotrimeric G-proteins are signal transducing complexes which are evolutionarily conserved in eukaryotes and play important roles in many different pathways. They consist of three subunits, Gα, Gβ and Gγ, and their basic mechanism of action has been elucidated in metazoans, where they are activated by membrane bound G-protein coupled receptors (GPCRs). The Gα subunit has GTPase activity and in the inactive, GDP-bound form associates with Gβ and Gγ, which form a stable heterodimer. Ligand binding of GPCRs leads to Gα nucleotide exchange, and the GTP-bound Gα dissociates from Gβγ. Subsequently, both Gα and the Gβγ dimer can relay signals to downstream targets. Signal transduction is terminated by the GTPase activity of Gα, which can be enhanced by GTPase Activity Promoting proteins (GAPs), and leads to re-association of the ternary complex (Oldham & Hamm, 2008; Pandey, 2019).

Plants possess canonical Gα, Gβ and Gγ subunits represented by GPA1, AGB1 and the redundantly acting AGG1/AGG2 in *Arabidopsis*. In addition, they contain plant-specific G-proteins, such as EXTRA LARGE G-PROTEINS (XLG1, 2 and 3 in *Arabidopsis*), a class of alternative Gα subunits with an N-terminal domain of unknown function. Similar to metazoans, signal transduction by plant heterotrimeric G-proteins is required for many processes, including growth, development, hormone and sugar sensing as well as responses to biotic and abiotic stresses (reviewed by Pandey, 2019; Zhong *et al*., 2019; Ofoe, 2021; Zhang *et al*., 2021). However, *bona fide* GPCRs have not been identified in plants and relatively little is known about the regulatory mechanisms. In contrast to animal heterotrimeric G-proteins, the canonical *Arabidopsis* Gα subunit GPA1 is inherently GTP-bound *in vitro* and likely constitutively active in plants. Signalling is thought to be controlled through deactivation via GAPs (Urano & Jones, 2014). For the Extra Large G-protein XLG2, only weak GTP binding / GTPase activity has been observed *in vitro* (Heo *et al*., 2012; Lou *et al*., 2020) and recent studies suggest that *in planta*, both GPA1 and XLG2 can fulfil many of their roles independently of nucleotide exchange (Maruta *et al*., 2019; Maruta *et al*., 2021).

Mounting experimental evidence indicates that plant heterotrimeric G-proteins act downstream of receptor-like kinases (RLKs) and are regulated by phosphorylation (Aranda-Sicilia *et al*., 2015; Roy Choudhury & Pandey, 2016; Tunc-Ozdemir *et al*., 2016; Pandey, 2020). A well-studied example is the complex consisting of XLG2, AGB1 and AGG1/AGG2 that transduces signals from pattern recognition receptors (PRRs). Liu *et al*. (2013) showed that *agb1* and *agg1 agg2* mutants, but not *gpa1* plants, have reduced immune responses to pathogen associated molecular patterns (PAMPs) such as flagellin, EF-Tu and chitin. XLG2 and XLG3 were identified as the Gα subunits that function in these defense pathways (Maruta *et al*., 2015). XLG2 was then shown to directly interact with the flagellin receptor complex components FLAGELLIN SENSING 2 (FLS2) and BOTRYTIS INDUCED KINASE 1 (BIK1) (Liang *et al*., 2016). The interaction with G-proteins stabilizes BIK1, presumably through inhibition of E3 ubiquitin ligases that target BIK1 for degradation (Wang *et al*., 2018). Consequently, *xlg2 xlg3* mutants contain less BIK1, leading to a decreased PAMP-induced ROS-burst by apoplastic NADPH oxidases. Upon flagellin treatment, XLG2 is phosphorylated in the N-terminal domain (possibly by BIK1) and dissociates from the receptor. Phosphorylation of XLG2 at four specific sites is required for full flagellin-induced ROS generation, even though this modification has no impact on BIK1 protein stability. Therefore, XLG2 is thought to modulate ROS generation via a second line of action, through direct interaction with the NADPH oxidase RBOHD (Liang *et al*., 2016). Moreover, XLG2 is known to interact with the E3 ubiquitin ligases PLANT U-BOX 2 (PUB2) and PUB4 (Wang *et al*., 2017), which are also required for a full PAMP-induced ROS-burst (Desaki *et al*., 2019; Derkacheva *et al*., 2020).

Further indications for an involvement of XLG2, AGB1 and AGG1/AGG2 in immune receptor signalling come from a suppressor screen performed with a mutant of the *BAK1 INTERACTING RECEPTOR-LIKE KINASE 1* (*BIR1*). BIR1 is a LRR-RLK that interacts with and probably negatively regulates BAK1 (BRI1-ASSOCIATED RECEPTOR KINASE 1). *bir1* knock-out mutants develop spontaneous cell death and show constitutive immune responses that are dependent on BAK1 and another LRR-RLK involved in immune signalling, SUPPRESSOR OF BIR1-1 (SOBIR1) (Gao *et al*., 2009; Liu *et al*., 2013; Liu *et al*., 2016). Mutations in *XLG2, AGB1* and *AGG1/AGG2*, but not *GPA1*, suppress the phenotype of *bir1* knockout plants (Liu *et al*., 2013; Maruta *et al*., 2015).

We previously identified and characterized *cerk1*-4, a mutant with an amino acid exchange in the extracellular domain of the *Arabidopsis* chitin receptor component CHITIN ELICITOR RECEPTOR KINASE 1 (CERK1). *cerk1*-4 plants show normal chitin signalling, but exhibit a hyperimmunity phenotype. Upon inoculation with powdery mildews, *cerk1-*4 mutants display excessive cell death formation and accumulation of the plant defence hormone salicylic acid (SA). Consequently, they do not support macroscopically visible growth of adapted powdery mildew species. The phenotype of *cerk1*-4 plants is associated with altered receptor processing and is conferred by the N-terminal part of the CERK1-4 protein. Unlike chitin signalling, manifestation of the *cerk1*-4 phenotype does not require CERK1 kinase activity (Petutschnig *et al*., 2014).

Here we show that *cerk1*-4 hyperimmunity is suppressed by mutations in *XLG2* and reduced in *cerk1*-4 *agb1* and *cerk1*-4 *agg1* plants. The formation of the *cerk1*-4 phenotype does not require BAK1 or SOBIR1 and is independent of NADPH-oxidase mediated ROS-generation. In depth cell biological and biochemical analyses of XLG2 mutant variants show that XLG2 fulfils its function in *cerk1*-4 signalling at the cell periphery. This novel role of XLG2 does not require its GTPase activity, but an intact N-terminal domain. Structural integrity of the N-terminal domain is also important for interaction with AGB1 and proper XLG2 phosphorylation. Finally, we show that the XLG2 interactor PUB2 contributes to *cerk1*-4 cell death.

## MATERIALS AND METHODS

### Plant growth conditions and pathogen infection assays

*Arabidopsis thaliana* (L.) Heynh. plants were grown under short day conditions (8h photoperiod) at 22°C (day) /18°C (night), 65% relative humidity and a light intensity of approximately 150 μmol m^−2^ s^−1^. Four-week-old plants were inoculated with either *Blumeria graminis* (DC.) Speer *f*.*sp. hordei* (*Bgh*) or *Erysiphe cruciferarum* Opiz ex L. Junell using a settling tower. Pictures and samples were taken at 5-10 dpi for *Bgh* and 7-14 dpi for *E. cruciferarum* infections.

### *cerk1-*4 suppressor screen and mapping of *les1*

Seeds of *cerk1-*4 (Petutschnig *et al*., 2014) were mutagenized with ethyl methanesulfonate (EMS) and resulting M2 plants were inoculated with either *Bgh* or *E. cruciferarum*. Plants that did not show any macroscopically visible cell death were selected and confirmed as suppressors in the M3 generation. For mapping, a strategy adapted from (Hartwig *et al*., 2012) was used. The *les1* mutant was backcrossed to *cerk1*-4 and approximately 250 F2 plants were scored for suppression of the powdery mildew-induced cell death phenotype. A pool of 31 clear suppressor plants and a pool of *cerk1*-4 plants grown in parallel were subjected to 100bp paired end Illumina sequencing. A coverage of approximately 40x was achieved for both samples. Using the CLC genomics workbench software, the reads were mapped to the Col-0 TAIR10 genome and SNPs with a frequency of 20% or higher were called. SNP tables were exported to Microsoft Excel, where SNPs present in *cerk1*-4 were subtracted from SNPs found in *les1*. For the remaining SNPs, the frequency was plotted against their chromosomal position (Fig. S1a). The obtained diagrams revealed a frequency peak on the lower arm of chromosome four, which contained a SNP leading to a premature stop codon (Q255*) in *XLG2* (*EXTRA-LARGE G-PROTEIN 2*, AT4G34390). Sequencing of *XLG2* in *les2* revealed an amino acid exchange (E293K) in the same protein. Both polymorphisms were confirmed as the causative mutations by complementation (Fig. S1c).

### Mutant analyses

The following previously characterized mutants were analysed and crossed with *cerk1*-4: *agb1*-2 (Ullah *et al*., 2003), *gpa1*-3 (SALK 066823) (Jones *et al*., 2003), *agg1*-1 *agg2*-1 (FLAG 197F06, SALK 010956) (Trusov *et al*., 2007), *xlg1*-2 (SALK 119657) (Maruta *et al*., 2015), *xlg2*-1 (SALK 062645) (Ding *et al*., 2008), *xlg3*-4 (SALK 107656) (Yu *et al*., 2018), *bak1*-4 (SALK 116202) (Kemmerling *et al*., 2007), *bak1*-5 (Schwessinger *et al*., 2011), *sobir1*-12 (SALK_050715) (Leslie *et al*., 2010), *sobir1*-14 (GABI_643F07) (Barghahn *et al*., 2021), *rbohD, rbohF, rbohD rbohF* (Torres *et al*., 2002), *pub2*-1 (SALK_086656) (Wang *et al*., 2017), *pub4*-3 (SAIL_859_H05) (Wang *et al*., 2013). *xlg1-*2, *xlg2*-1 and *xlg3*-4 were used to produce double and triple mutants.

### Quantitative RT-PCR and ROS-burst assays

Quantitative RT-PCR was performed as described previously (Petutschnig et al., 2014). For ROS burst assays, 4 mm leaf discs were cut out and floated in water in a white 96-well plate for at least 4 h. Then the water was replaced with a solution containing 1μg/ml horse radish peroxidase (Sigma/Merck), 50μM L-012 (Wako Chemicals) and 100nM flg22. Chemiluminescence was immediately recorded using a Tecan Infinite M200 plate reader. Statistical significance was tested with unpaired, two-tailed t-tests.

### Protein Work

Protein extraction from *Arabidopsis* leaves and Western blotting were performed according to previously published protocols (Petutschnig *et al*., 2010). The antibodies used in Western blots were anti-GFP (Chromotek), anti-FLAG M2 (Sigma), anti-CERK1 (Petutschnig *et al*., 2010), anti-GPA1 (Agrisera), anti-AGB1 (Agrisera) and anti-XLG2, a custom polyclonal antibody raised against the first 200 amino acids of XLG2 (Agrisera). Western Blot Signal Enhancer (Pierce / Thermo Fisher Scientific) was used for development with GFP and XLG2 antibodies. For GFP pull-downs, standard protein extracts were prepared (Petutschnig *et al*., 2010) and the target protein was isolated with either Chromotek GFP trap beads (mass spectrometry experiments) or Nanotag GFP selector beads (Co-IP experiments). Protein extracts were mixed with αGFP nanobody beads at a ratio of approximately 0.2-0.5mg total protein / μl bead slurry and incubated at 4°C on a rotating wheel overnight. The next day, the beads were pelleted by gentle centrifugation at 1000 × g or slower. The supernatant was discarded and the beads were washed 2-3 times with extraction buffer (Petutschnig *et al*., 2010) and 2-3 times with TBS-T. Bound proteins were released by boiling beads in 1x SDS loading dye or beads were directly used for trypsin digestion.

### Microscopy

Confocal microscopy was performed on a Leica TCS SP8 Falcon system equipped with HyD SMD detectors, a pulsed white light laser and the LASX 3.5.5 software. Venus fluorescent protein was excited at 514nm and fluorescence was detected between 525-560nm. Since *pXLG2-Venus-XLG2* and derivatives are weakly expressed, the accumulation mode was used in some experiments. A fluorescence lifetime gate was set between 0.4-6ns to exclude chloroplast autofluorescence signals from images. Excitation and detection wavelengths for subcellular marker proteins mKate2-Lti6b and mTurquoise2-N7 (Ghareeb *et al*., 2016) were 594nm/600-641nm and 458nm/462-485nm respectively. Stitched images were recorded and assembled with the LASX Navigator module. Z-stacks were exported as maximum projections and brightness/contrast was adjusted either with Adobe Photoshop CS5 or ImageJ (Fiji distribution).

## RESULTS

### A suppressor screen identifies XLG2 as an essential component of the *cerk1-*4 cell death pathway

In order to identify molecular components involved in the establishment of CERK1-4-dependent hyperimmunity, we performed a genetic suppressor screen. For this, we infected an EMS-mutagenized population of *cerk1-*4 with powdery mildews and looked for plants with wild type-like interaction phenotypes. Two recessive suppressor mutations fully inhibited the development of dysregulated cell death upon inoculation with the incompatible powdery mildew fungus *Blumeria graminis f. sp. hordei* (*Bgh*) (Fig. 1a) and the compatible *Erysiphe cruciferarum* (Fig. 1b). Therefore, they were named *<u>le</u>sion <u>s</u>uppressor 1* and *2* (*les1* and *les2*). Neither one of the mutations led to any obvious developmental changes in non-inoculated plants (Fig. 1c). *Bgh*-induced *PATHOGENESIS RELATED1* (*PR1*) expression, which is increased in *cerk1*-4, was reduced to levels similar to the wild type in the *les1* and *les2* mutants (Fig. 1d).

**Figure 1:**
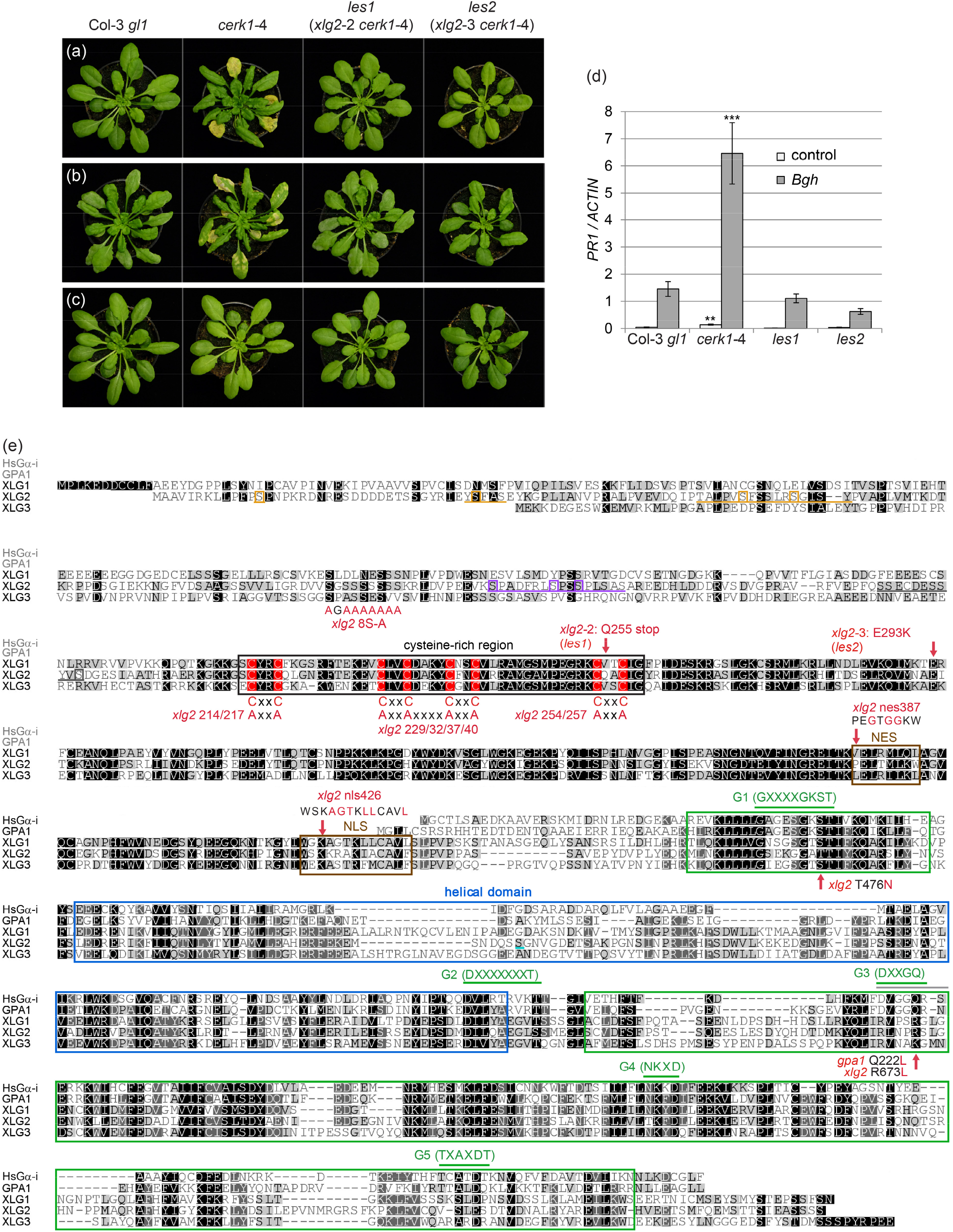
Two independent mutations in *XLG2* suppress enhanced powdery mildew-induced defence responses in *cerk1-*4. Representative plants of the indicated genotypes 10 days after inoculation with (a) *Blumeria graminis f. sp. hordei (Bgh)* or (b) *Erysiphe cruciferarum*. (c) Unchallenged control plants of the same age. (d): Quantitative RT-PCR analysis of *PR1* expression in plants infected with *Bgh* (5dpi) normalized to the expression of *ACTIN*. The results represent averages of four technical replicates ± SD. Significant differences between mutants and Col-3 *gl1*: ** P < 0.01, *** P < 0.001. (e) Alignment of Arabidopsis XLG proteins with GPA1 and human Gα-i. Within the GTPase domain, the helical and Ras-like domains (as determined for GPA1) are shown as blue and green boxes, respectively (Jones *et al*., 2011). The conserved GTPase regions G1-G5 are indicated by green lines and consensus sequences for motifs are given in brackets (Ding *et al*., 2008). The cysteine rich region is depicted as a black box and conserved cysteines within are highlighted in red. A functional nuclear export signal (NES) determined for XLG3 and nuclear localisation signal (NLS) determined for XLG2 (Chakravorty *et al*., 2015) are indicated by brown boxes. Mutations generated in this study are indicated in red. Regions within the N-terminal domain that show reduced phosphorylation in XLG2^E293K^ compared to wild type XLG2 are underlined in orange, a region with increased phosphorylation in XLG2^E293K^ is underlined purple and a region with unchanged phosphorylation levels is underlined grey. Unambiguously assigned phosphorylation sites within these regions are indicated by boxes. The phosphorylation sites within the purple region were shown to be required for full flg22-induced ROS-burst (Liang *et al*., 2016).

We identified the *les1* mutation using a mapping-by-sequencing approach (Hartwig *et al*., 2012). Single nucleotide polymorphism (SNP) linkage analysis located the *les1* mutation at the lower arm of chromosome 4 (Fig. S1a), with a mutation introducing a premature stop codon (Q255*) in *XLG2* (Fig. 1e) being the most promising candidate. Interestingly, we found that our second suppressor mutant, *les2*, also harbours a mutation in the *XLG2* gene, which results in a glutamic acid to lysine exchange (E293K) within a highly conserved region of the XLG2 N-terminal domain (Fig. 1e). Consequently, we transformed both mutants with a construct containing the wild type *XLG2* promoter and coding region, and found that the *cerk1-*4 cell death response to powdery mildew infection was restored in *les1* as well as *les2* (Fig. S1b and c). Together, these complementation analyses suggest that *cerk1*-4-associated hyperimmunity requires functional XLG2. Since both suppressor mutations compromise XLG2, we renamed the suppressor loci in *les1* and *les2* to *xlg2*-2 and *xlg2*-3, respectively.

### Mutations in Gα and Gβ subunits AGB1 and AGG1 partially suppress the *cerk1-*4 phenotype\

To investigate the role of other *Arabidopsis* heterotrimeric G-proteins in *cerk1-*4 cell death control, we crossed *cerk1-*4 with mutants of canonical Gα, Gβ and Gγ subunits as well as all three XLGs. The resulting double mutants were analysed for their reaction to *Bgh*. As expected, the *xlg2*-1 T-DNA insertion fully suppressed the *cerk1*-4 cell death response and restored expression of the SA marker *PR1* to wild type levels (Fig. 2a,b). In contrast, mutations in *GPA1, XLG1* and *XLG3* did not suppress *cerk1-*4. Notably, both *agb1*-2 *cerk1-*4 and *agg1*-1 *cerk1-*4 double mutants showed an intermediate phenotype concerning cell death formation as well as *PR1* expression (Fig. 2a,b). In *agg1*-1 *cerk1-*4, this might be due to functional redundancy between AGG1 and AGG2, since generation of a *cerk1-*4 *agg1-*1 *agg2-*1 triple mutant failed because of genetic linkage between *CERK1* (At3g21630) and *AGG2* (At3g22942). *AGB1*, however, is a single copy gene in *Arabidopsis*. Thus, we performed Western blotting with specific antibodies to test if CERK1 or XLG2 abundance is reduced in the G-protein mutants. CERK1 levels in these mutants were comparable to wild type (Fig. 2c), but interestingly, XLG2 had a slightly lower apparent mass in *agb1* and *agg1*-1 *agg2*-1 (Fig. 2c). These findings suggest that XLG2 might require protein modifications for its function that are (partially) dependent on AGB1 and AGG1/2. GPA1 and AGB1 Western blots were included in the experiment as controls and revealed drastically reduced AGB1 abundance in *agg1*-1 *agg2*-1, probably because βγ-dimer formation is a prerequisite for AGB1 stability (Fig. 2c).

**Figure 2:**
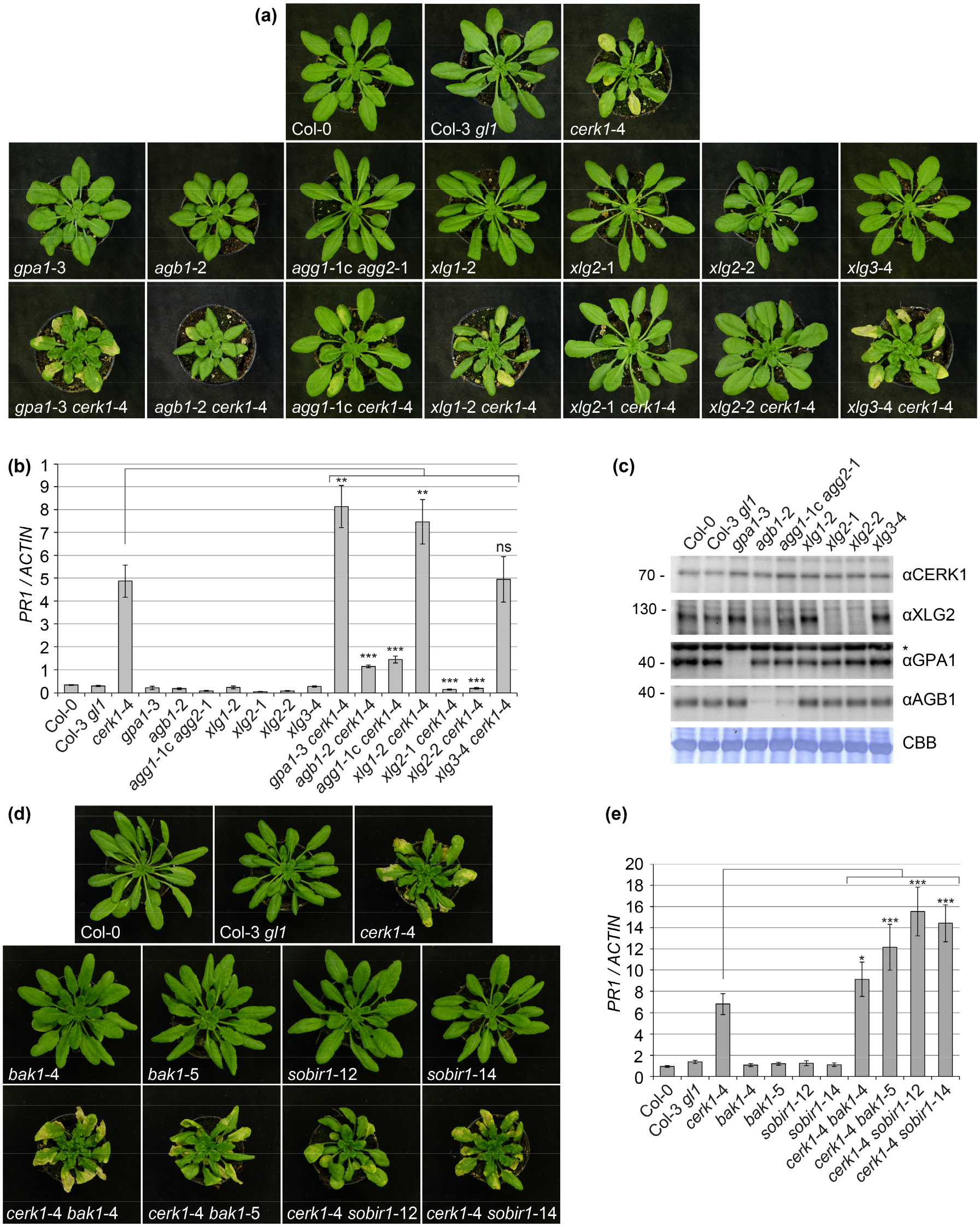
G-protein β and γ subunits, but not BAK1 and SOBIR1, are required for full development of the *cerk1-*4 cell death phenotype. **(a)** The indicated heterotrimeric G-protein single and *cerk1-*4 double mutants were inoculated with *Bgh* and images were taken 7 dpi. **(b)** *PR1* expression was analysed by qRT-PCR and values were normalized to the expression of *ACTIN*. Whole rosettes were harvested 7d after *Bgh* inoculation and 10 plants were pooled for each sample. **(c)** G-protein mutants were analysed by Western blotting using *CERK1-*, GPA1-, AGB1- and XLG2-specific antibodies. * indicates unspecific band. CBB: Coomassie Brilliant Blue staining. **(d)** The indicated *bak1* and *sobir1* mutants and their crosses with *cerk1-*4 were inoculated with *Bgh* and images were taken 7dpi. **(e)** *PR1* expression was analysed by qRT-PCR and values were normalized to the expression of *ACTIN*. Whole rosettes were harvested after 5d and 10 plants were pooled for each sample. qRT-PCR experiments are presented as averages of four technical replicates ± SD. Significant differences between *cerk1*-4 and *cerk1*-4 double mutants are indicated, * P < 0.05, ** P < 0.01, *** P < 0.005.

### Cell death signalling in *cerk1*-4 is not mediated by BAK1 or SOBIR1-containing receptor complexes

Hyperimmunity phenotypes of the *cerk1*-4 mutant are associated with the extracellular part of the CERK1-4 protein (Petutschnig *et al*., 2014) and could potentially be perceived by a yet unknown RLK that signals through XLG2. Many of the RLKs and RLPs characterized to date use BAK1 as a co-receptor and/or SOBIR1 as an adaptor kinase (Escocard de Azevedo Manhaes *et al*., 2021). This encouraged us to test if the *cerk1*-4 cell death phenotype is dependent on BAK1 and SOBIR1. Phenotypic analysis of powdery mildew-infected *cerk1*-4 *bak1* and *cerk1*-4 *sobir1* double mutants clearly demonstrated that BAK1 and SOBIR1 are dispensable (Fig. 2d,e). *PR1* expression appeared to be even higher in *cerk1*-4 *bak1* and *cerk1*-4 *sobir1* compared to *cerk1-*4 plants (Fig. 2e).

### The role of XLG2 in *cerk1-*4 signalling is independent of its function in PAMP-induced ROS generation

XLG2 is required for full flagellin-induced ROS generation. It associates with the flagellin receptor complex and regulates NADPH oxidase activity through multiple modes of action (Liang *et al*., 2016; Liang *et al*., 2018; Wang *et al*., 2018). Similar to previous reports (Maruta *et al*., 2015), we observed a moderate reduction of flg22-induced ROS in the *xlg2*-1 mutant, but robustly reduced ROS generation in *xlg2 xlg3* double and *xlg1 xlg2 xlg3* triple mutants (Fig. 3a). This redundant action of XLG proteins argues against a role for ROS generation in *cerk1*-4 hyperimmunity, which can be suppressed by mutations in XLG2 alone. For further analysis, we made use of a mutated XLG2 variant lacking four *in vivo* phosphorylation sites (XLG2^S141A, S148A, S150A, S151A^, further referred to as XLG2^4S-A^) required for full flg22-induced ROS burst (Liang *et al*., 2016). We introduced XLG2^4S-A^-FLAG into the *xlg2-*2 *cerk1-*4 double mutant. Wild type XLG2-FLAG and the phosphomimic version XLG2^4S-D^-FLAG (Liang *et al*., 2016) were included as controls. All three XLG2-FLAG versions were similarly able to re-establish the *cerk1-*4 phenotype, indicating that the XLG2 phosphorylation sites governing flg22-dependent ROS regulation do not play a role here (Fig. 3b,c, Table S2). To corroborate these findings, we crossed *cerk1-*4 with the NADPH oxidase mutants *rbohD* and *rbohF* and the respective double mutant (Torres *et al*., 2002). Neither loss of RbohD or RbohF, nor the combined loss of both, suppressed the *cerk1*-4 cell death phenotype upon powdery mildew infection (Fig. 3d,e). Taken together, these data indicate that XLG2 exerts a function in *cerk1*-4 hyperimmunity that is independent of NADPH oxidase-generated ROS.

**Figure 3:**
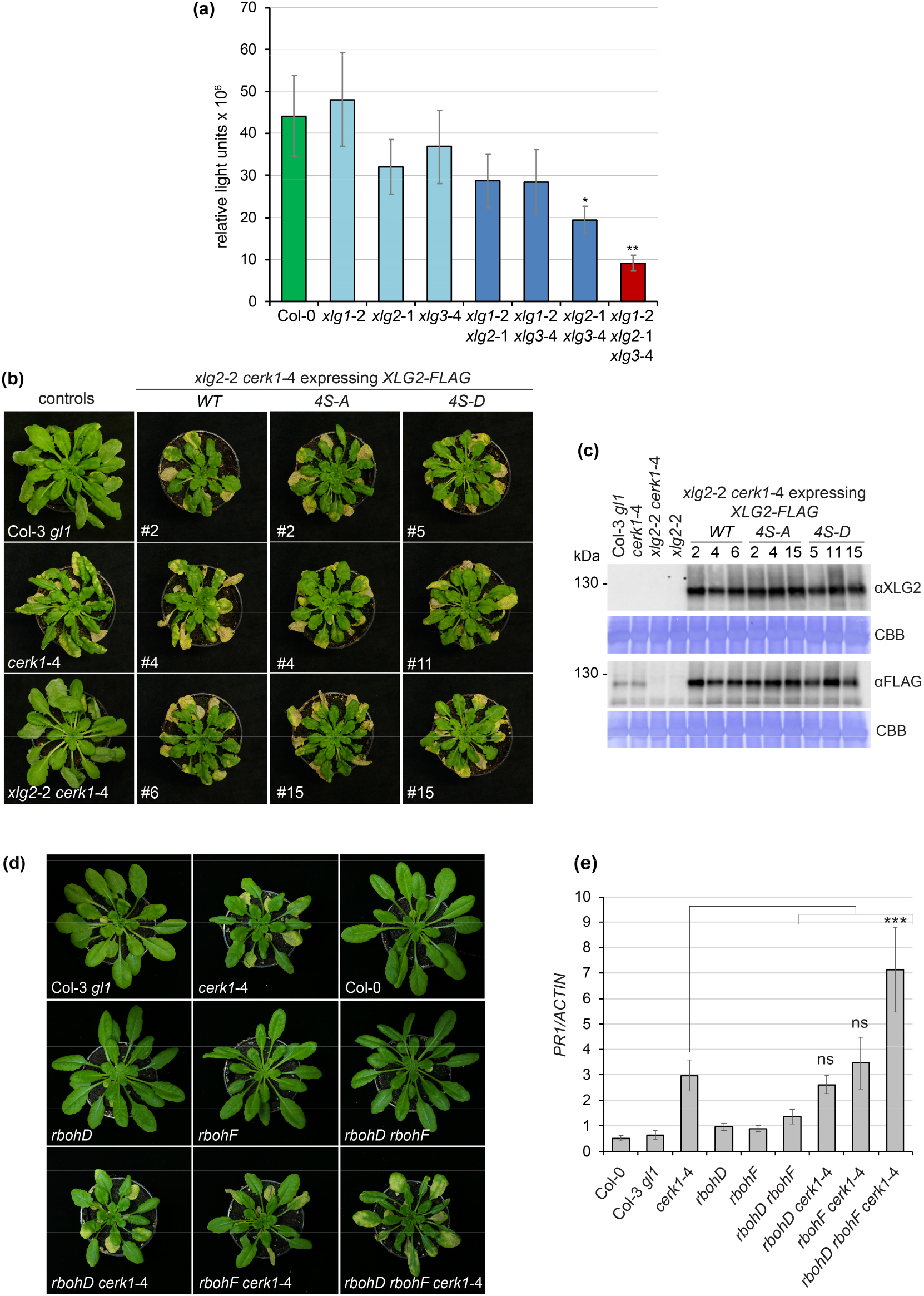
The *cerk1*-4 cell death phenotype is independent of NADPH oxidase-mediated ROS generation. **(a)** The indicated plant lines were treated with 100 nM flg22 and ROS generation was recorded over 50 min. Data represent average accumulated values from 6 experiments (with 8 samples per line and experiment) ± SEM. * P < 0.05, ** P < 0.01. **(b)** *xlg2*-2 *cerk1-*4 plants were transformed with the indicated XLG2-FLAG constructs. 10 days after inoculation with *E. cruciferarum*, all transgenic plants showed cell death symptoms similar to *cerk1-*4. Three representative transformants per construct are shown. **(c)** Western blots of the plants shown in (B) with FLAG- and XLG2-specific antibodies. **(d, e)** *cerk1-*4 plants were crossed with *rbohD* and *rbohF*. The resulting double and triple mutants and control plants were inoculated with *Bgh* and analysed 7 dpi. **(d)** Pictures of representative plants. **(e)** For each line, 10 whole rosettes were pooled, PR1 expression was analysed by qRT-PCR and values were normalized to the expression of *ACTIN*. qRT-PCR experiments are presented as averages of four technical replicates ± SD. Significant differences between *cerk1*-4 and *cerk1*-4 crosses are indicated, *** P < 0.005, (ns, not significant).

### Cell periphery localisation of XLG2 is required for *cerk1*-4 cell death dysregulation

When transiently expressed in *N. benthamiana* leaves, XLG2 localises to the cell periphery as well as the nucleus (Chakravorty *et al*., 2015; Maruta *et al*., 2015). We could confirm this finding with Venus-tagged XLG2 (Fig. S2a) and observed the same localisation upon particle bombardment in *Arabidopsis* epidermal cells (Fig. S2b). Interestingly, the non-functional XLG2^E293K^ variant that we identified in the *les2*/*xlg2*-3 mutant almost exclusively localised to the nucleus in both transient systems (Fig. S2a,b). To study the subcellular localisation and its importance for XLG2 function in more detail, we analysed Venus-XLG2 and Venus-XLG2^E293K^ expressed under the control of the *XLG2* promoter in stably transformed *xlg2*-2 and *cerk1*-4 *xlg2*-2 plants. In these, Venus-XLG2 localised predominantly to the cell periphery and Venus-labelled nuclei were rarely found (Fig. 4a, Table S3, S4), suggesting that nuclear accumulation of XLG2 observed in transient systems may be a stress response caused by *Agrobacterium* infiltration or particle bombardment. Again, XLG2^E293K^ showed almost exclusive localisation to nuclei (Fig. 4a, Table S3, S4). Localisation patterns were very similar in the *CERK1* and *cerk1*-4 background. Since dysregulated cell death responses of *cerk1*-4 plants become apparent upon infection with powdery mildews, we investigated the subcellular behaviour of Venus-XLG2 and XLG2^E293K^ after inoculation with *E. cruciferarum*. Fungal infection caused increased accumulation of Venus-XLG2 in the attacked epidermal cells, while distant cells were not affected. Moreover, nuclear Venus-XLG2 signals could be detected at such interaction sites (Fig. 4b, Fig. S2c). XLG2^E293K^ also showed increased abundance in cells challenged with *E. cruciferarum*, but retained its predominantly nuclear localisation (Fig. 4b, Fig. S2c). Notably, Venus-XLG2 was able to re-establish the *cerk1*-4 phenotype in *xlg2-*2 *cerk1-*4 plants, whereas the mutant variant Venus-XLG2^E293K^ was not (Fig. 4c, Table S4). Western blotting was employed to confirm accumulation of full-length fusion proteins in the transgenic lines (Fig. 4d). These experiments revealed that Venus-XLG2^E293K^ migrated faster in SDS-PAGE than the wild type fusion protein, which will be discussed in more detail further below.

**Figure 4:**
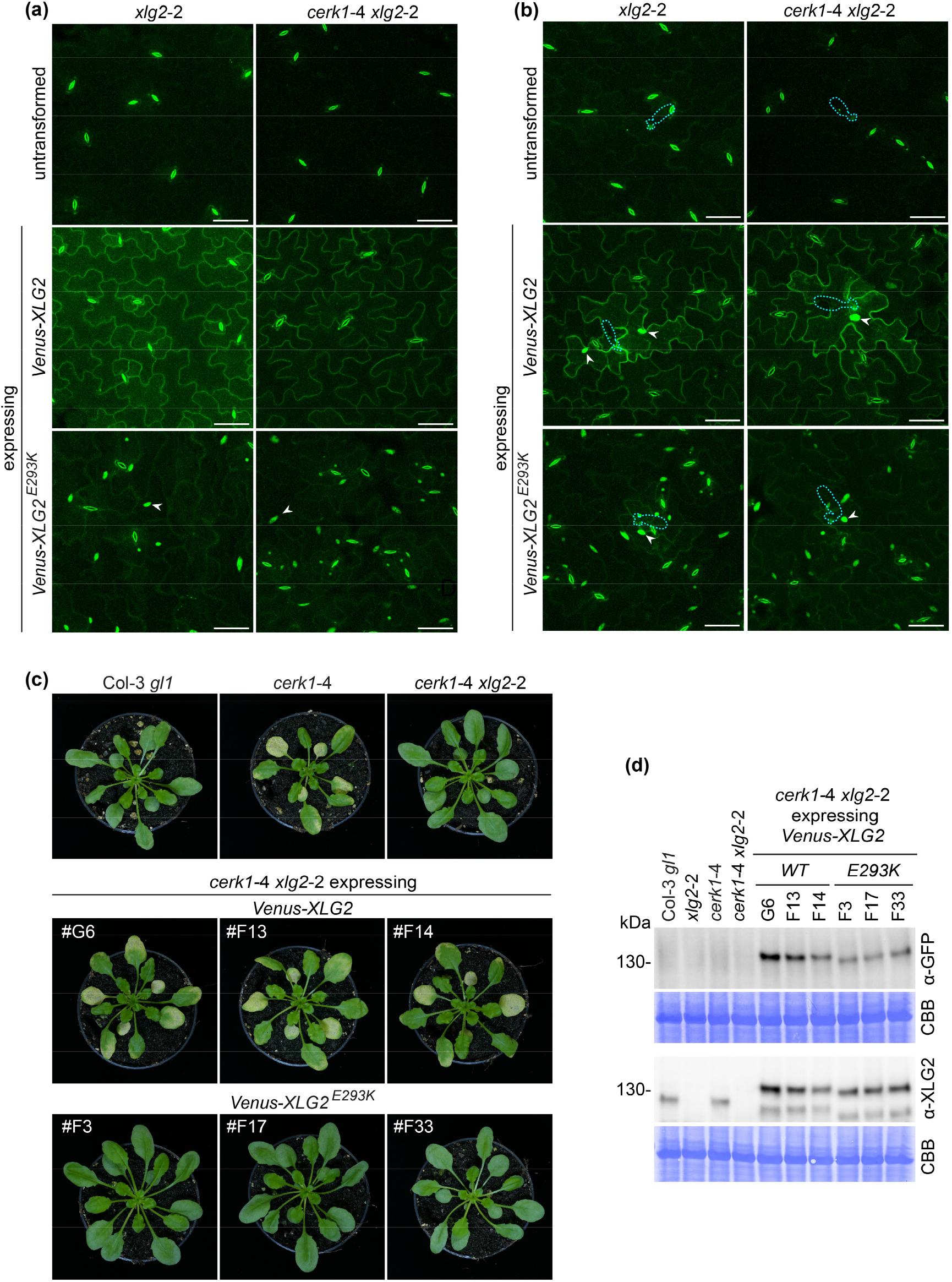
The E293K mutation alters subcellular localisation of XLG2 in unchallenged and powdery mildew infected *Arabidopsis* plants and impairs XLG2 function. (a, b) Localisation analyses of Venus-XLG2 and Venus XLG2^E293K^ stably expressed from the endogenous promoter in *xlg2*-2 and *cerk1*-4 *xlg2*-2. Untransformed control plants are included for better discernibility of stomata autofluorescence. Leaf epidermis CLSM images of (a) unchallenged plants, (b) plants 1 dpi with *E. cruciferarum*. Images are maximum projections of z-stacks spanning the epidermal cell layer. Positions of spores and appressoria are indicated by dashed blue outlines and example nuclei are indicated by white arrowheads. Scale bar = 50 μm. (c) *cerk1*-4 *xlg2*-2 plants stably expressing *pXLG2-Venus-XLG2* or *Venus-XLG2*^*E293K*^ as well as the indicated controls 10 dpi with *E. cruciferarum*. Images of three independent transformants are shown. (d) Western Blots of the transgenic *pXLG2-Venus-XLG2* or *Venus-XLG2*^*E293K*^ lines developed with XLG2 and GFP-specific antibodies. CBB = Coomassie Brilliant Blue stained membrane.

Since the non-functional mutant variant XLG2^E293K^ accumulated predominantly in the nucleus, we speculated that the cell periphery localisation of wild type XLG2 might be essential for its role in *cerk1*-4 cell death signalling. To address this question, we attached nuclear localisation and export signals (NLS and NES) (Garcia *et al*., 2010) to the C-terminus of Venus-XLG2. Non-functional sequences (nls and nes) were included as controls. Venus-XLG2-NLS showed nuclear localisation without pathogen challenge, but still had considerable signals at the cell periphery. Importantly, Venus-XLG2-NES was found only at the cell periphery in unchallenged tissues and did not show the typical stained nuclei in powdery mildew infected cells (Fig. 5a). All variants accumulated to similar protein levels and were able to complement *xlg2-*2 *cerk1-*4 (Fig. 5b,c, Table S4). The data suggest that indeed the periphery-localised fraction of the XLG2 protein pool promotes *cerk1*-4 dependent cell death and that nuclear localisation is dispensable for this process.

**Figure 5:**
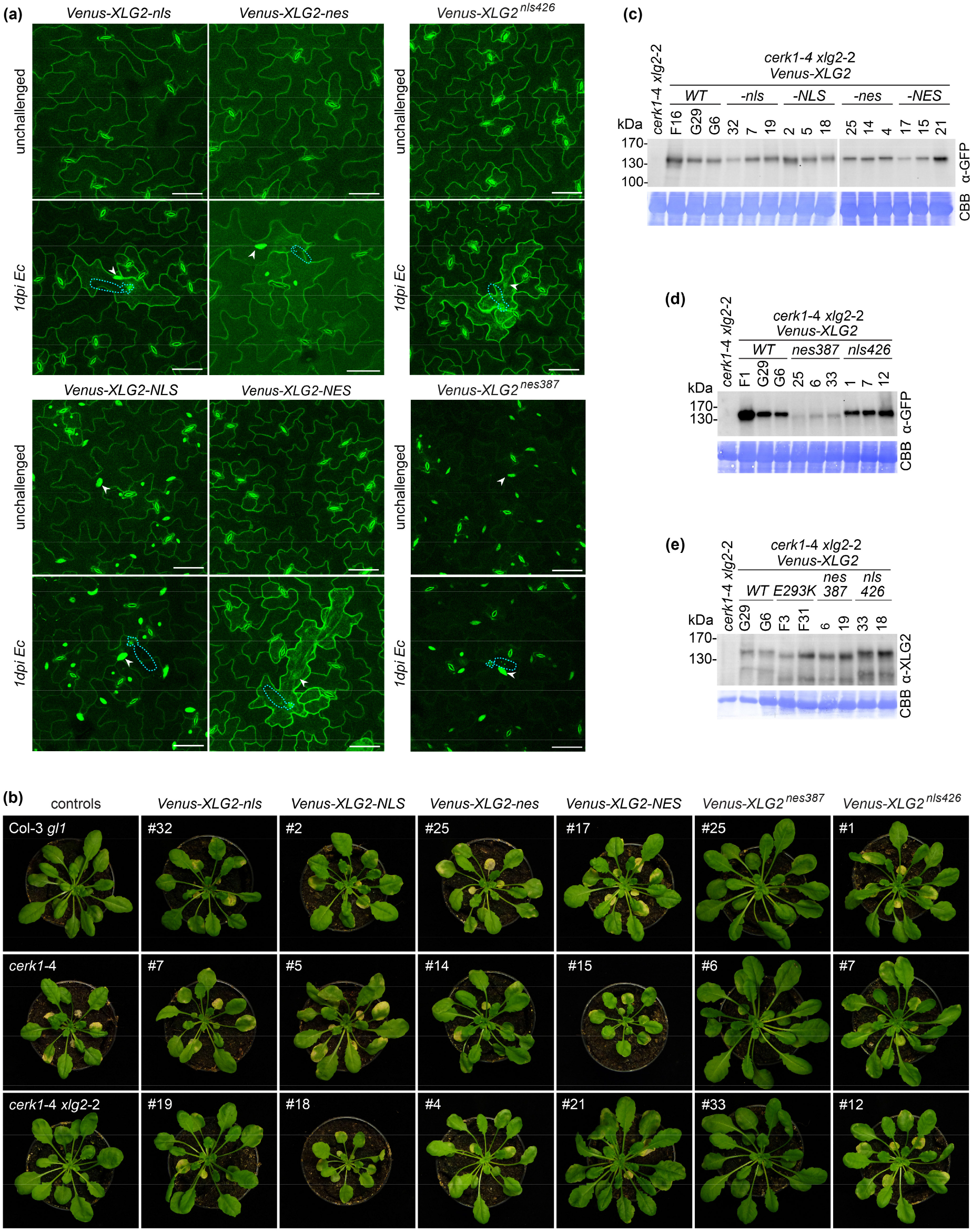
Cell periphery localisation of XLG2 is required for *cerk1*-4 cell death signalling. *cerk1*-4 *xlg2*-2 plants were transformed with *pXLG2*-*Venus-XLG2-*nls/NLS/nes/NES and with *pXLG2*-*Venus-XLG2* containing nes387 or nls426 mutations. (a) CLSM analysis of unchallenged leaves and leaves infected with *E. cruciferarum* (1 dpi). Images are maximum projections of z-stacks spanning the epidermal cell layer. Example nuclei are marked by arrowheads and the positions of fungal spores at sites of attempted penetration are outlined by dashed blue lines. Scale bar = 50 μm. (b) The indicated transgenic lines and controls were inoculated with *E. cruciferarum* and pictures were taken 14 dpi. Three independent lines per construct are shown. Transgenic protein levels in (c) Venus-XLG2-nls/NLS/nes/NES lines and (d) Venus-XLG2^nes387^/ XLG2^nls426^ lines were confirmed by Western blotting after *E. cruciferarum* infection. (e) The apparent mass of Venus-XLG2^nes387^ and Venus-XLG2^nls426^ was compared to Venus-XLG2 WT and Venus XLG2^E293K^ in a side-by-side Western Blot. CBB = Coomassie Brilliant Blue stained membrane.

### Intrinsic features of the N-terminal domain control XLG2 subcellular localisation and *cerk1*-4 cell death signalling

In order to analyse the regulation of XLG2 subcellular localisation, we mutated an internal NLS (nls426) and NES (nes387) within the XLG2 N-terminal domain (Fig. 1e) based on a study by Chakravorty *et al*. (2015). Stable transgenic expression from the *XLG2* promoter showed that Venus-XLG2^nls426^ localised to the cell periphery, had wild-type like migration properties in Western blots and could complement *xlg2-*2 *cerk1-*4. In contrast, Venus-XLG2^nes387^ was exclusively found in the nucleus, migrated faster in SDS-PAGE and was non-functional in *cerk1*-4 cell death signalling (Fig. 5a,b,d,e, Table S4). Taken together, the data corroborated that XLG2’s specific role in *cerk1*-4 mutants is fulfilled at the cell periphery, and indicated that subcellular localisation of XLG2 is determined by sequences within the N-terminal domain of the protein. N-terminal domains of XLG proteins contain highly conserved cysteine rich regions with four invariant CxxC motifs (Fig. 1e), which are possible zinc finger domains (Lee & Assmann, 1999). Mutation of one (C214/217A and C254/257A) or two (C229/232/237/240A) CxxC motifs (Fig.1e) resulted in non-functional Venus-XLG2 fusion proteins that resembled Venus-XLG2^E293K^ and XLG2^nes387^ in confocal analyses and Western blots (Fig. S3a-d, Table S4). Thus, correct localisation of XLG2 also requires structural integrity of the cysteine rich region.

XLG proteins harbour a Gα domain and weak GTP binding and GTPase activity have been demonstrated for XLG2 *in vitro* (Heo *et al*., 2012; Lou *et al*., 2020). To investigate if XLG2 GTPase activity plays any role in formation of the *cerk1*-4 cell death phenotype, we mutated its GTP binding site (T476N) as described previously (Heo *et al*., 2012; Maruta *et al*., 2021) (Fig. 1e). In addition, we introduced a mutation that could potentially affect GTPase activity (R673L), since a mutation in this position renders GPA1 inactive (Okamoto *et al*., 2001; Ullah *et al*., 2003)(Fig. 1e). Subcellular localisation and running behaviour in Western blots were unaffected in XLG2^T476N^ as well as XLG2^R673L^ and both mutant proteins were able to complement *xlg2*-2 *cerk1-*4 (Fig. S4, Table S4). This indicates that GTP-binding and hydrolysis are irrelevant for XLG2 positioning and activity in *cerk1*-4 cell death signalling.

### Gβγ modulates protein modifications and subcellular localisation of XLG2

Notably, the SDS-PAGE mobility shift of Venus-XLG2^E293K^ (Fig. 4d), XLG2^nes387^ (Fig. 5d,e) and the cysteine rich region mutants (Fig. S3c,d) resembled the behaviour of wild type XLG2 in *agb1* and *agg1 agg2* mutants (Fig. 2c). However, direct comparison of *xlg2-*3 (containing untagged XLG2^E293K^), *agb1* and *agg1 agg2* showed that the XLG2 band shift in *xlg2-*3 is more pronounced than in the Gβγ mutants (Fig. 6a). We speculated that XLG2 might carry post-translational modifications, which are moderately reduced in *agb1* and *agg1 agg2*, but more strongly affected or missing in XLG2^E293K^. To investigate this hypothesis, we transformed *agb1*-2 mutants with *pXLG2-Venus-XLG2* (WT) as well as *pXLG2-Venus-XLG2*^*E293K*^ (Table S5). Western blots showed that Venus-XLG2 migrated at a lower apparent mass in the AGB1-deficient background (Fig. 6b). Venus-XLG2^E293K^ however showed no such mobility shift (Fig. 6c), suggesting that the same type of modification is affected in *agb1*-2 and the XLG2^E293K^ mutant. In agreement with this notion, Venus-XLG2 localised to both, nucleus and cell periphery in unchallenged *agb1*-2 plants, while Venus-XLG2^E293K^ localisation was not affected (Fig. 6d). XLG2 is known to interact with the Gβγ dimer (Zhu *et al*., 2009; Chakravorty *et al*., 2015; Maruta *et al*., 2015). However, this interaction capacity was strongly impaired (with and without PAMP treatment) in the XLG2^E293K^ variant (Fig. 6e). These data favour a model where direct interaction of XLG2 with Gβγ mediates full post-translational XLG2 modification and contributes to proper XLG2 subcellular localisation.

**Figure 6:**
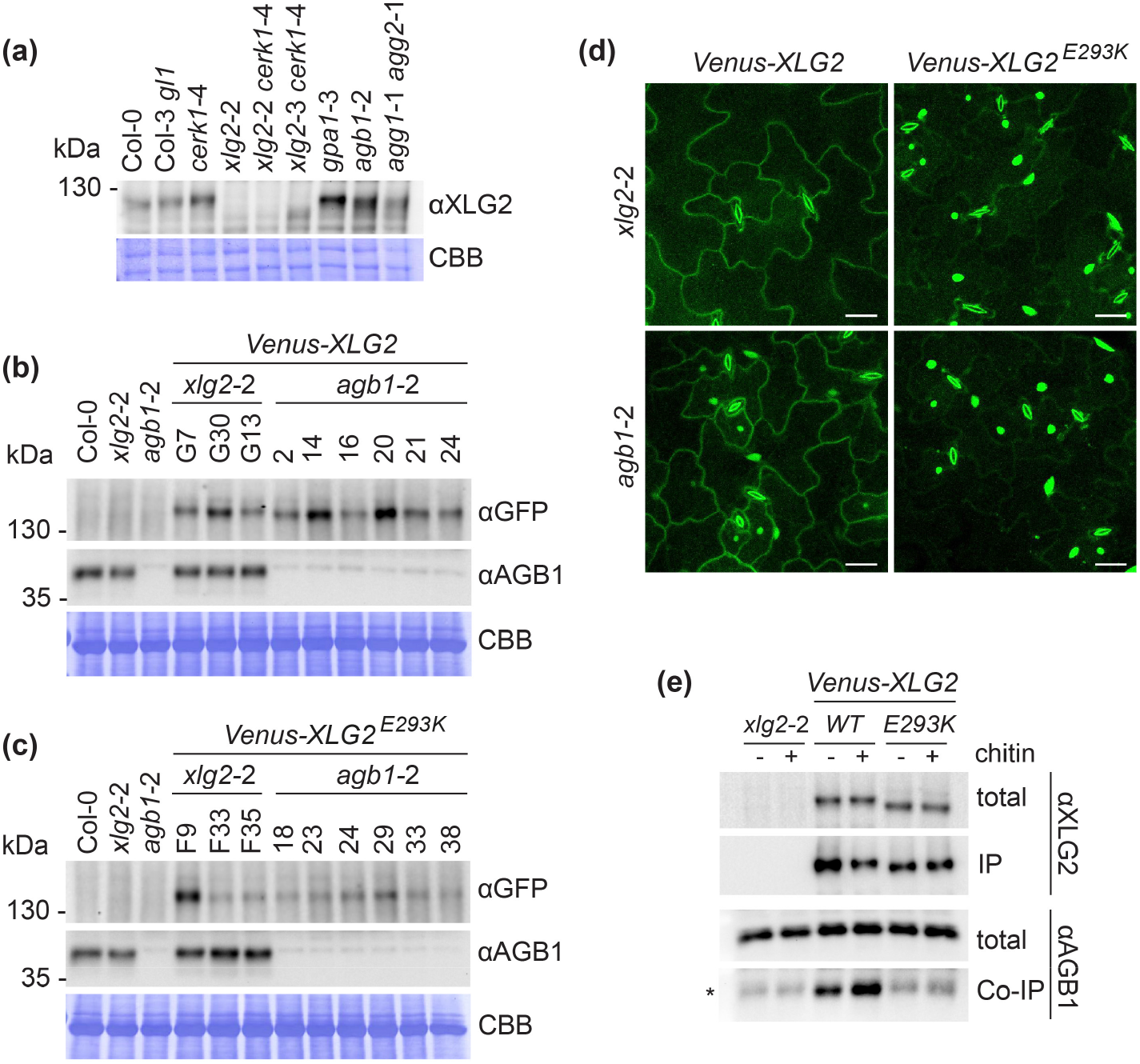
XLG2 localisation and protein modification are altered in *agb1*-2 and the non-functional XLG2^E293K^ variant. (a) A Western blot of the indicated *Arabidopsis* lines with a specific XLG2 antibody reveals band shifts in *xlg2*-3 (XLG2^E293K^), *agb1* and *agg1 agg2*. (b) *Venus-XLG2 WT* and (c) *Venus XLG2*^*E293K*^ were stably expressed in *xlg2-*2 and *agb1*-2. Western blots were performed with antibodies against Venus (αGFP) and AGB1. CBB: Coomassie brilliant blue stained membranes. (d) CSLM images of Venus-XLG2 and Venus-XLG2^E293K^ stably expressed in *xlg2*-2 and *agb1*-2. Images are maximum projections of 12 focal planes recorded 1 μm apart. Size bar: 25 μm. (e) Venus-XLG2/XLG2^E293K^– AGB1 Co-IP assay. Total protein extracts (input) were prepared from untransformed *xlg2*-2 as well as *xlg2*-2 stably expressing Venus-XLG2 WT or Venus-XLG2^E293K^. Pull-downs were performed with nanobody beads recognizing GFP/Venus. Input and pull down fractions were analysed using αGFP/Venus and αAGB1 antibodies. *AGB1 shows low levels of unspecific binding to the beads in absence of XLG2.

### Phosphorylation in the N-terminal region of XLG2 is not essential for *cerk1*-4 cell death signalling

Since XLG2 is heavily phosphorylated in the N-terminal domain (Liang *et al*., 2016; Chakravorty & Assmann, 2018) we purified Venus-XLG2 and Venus-XLG2^E293K^ from transgenic *Arabidopsis* plants via the fluorescence tag and performed phosphatase assays. Upon de-phosphorylation, both XLG2 versions showed the same running behaviour in Western blot analyses (Fig. 7a), indicating that Venus-XLG2^E293K^ indeed has a phosphorylation defect. For further analysis, we carried out phosphoproteomics studies on the two fusion proteins, each isolated from three independent *Arabidopsis* lines. Phosphopeptides were clustered into 5 non-overlapping regions (Fig. 1e, Fig. 7b), and subjected to targeted selected ion monitoring (tSIM) to quantify phosphorylation levels. These analyses revealed that XLG2^E293K^ showed drastically reduced phosphorylation near the N-terminus (aa 1-79, Fig. 7b). Interestingly, phosphorylation of sites reported to regulate XLG2 function in PAMP-induced ROS generation (aa 141-156) (Liang *et al*., 2016) was increased in XLG2^E293K^, while more distant sites (aa 184-194) were not affected (Fig. 7b). We had already shown that phosphorylation within the aa 141-156 region does not play a role in *cerk1*-4 signalling (Fig. 3b,c). In order to assess the contribution of XLG2 N-terminal phosphorylation, we mutated all 27 serine and threonine residues in the aa 1-79 segment to alanines (XLG2^27S/T-A^). XLG2 has a serine rich stretch (SGS_7_ at amino acid positions 124-130, Fig. 1e), which is not amenable to mass spectrometry analysis, but may still be important for XLG2 function. This was also mutated (XLG2^8S-A^) and both constructs were transformed into *xlg2*-2 *cerk1-*4. The expressed fusion proteins showed reduced apparent masses compared to the wild type Venus-XLG2 version (Fig. 7c). Venus-XLG2^27S/T-A^ displayed a very similar motility in SDS-PAGE to Venus-XLG2^E293K^ (Fig. 7d). However, both Venus-XLG2^8S-A^ and Venus-XLG2^27S/T-A^ showed a cell periphery localisation comparable to the wild type version (Fig. 7e) and complemented the *xlg2*-2 mutation (Fig. 7f). These data suggest that the mobility shift of XLG2^E293K^ is due to reduced phosphorylation, but this is not the cause of its altered localisation and inability to signal in the *cerk1*-4 pathway.

**Figure 7:**
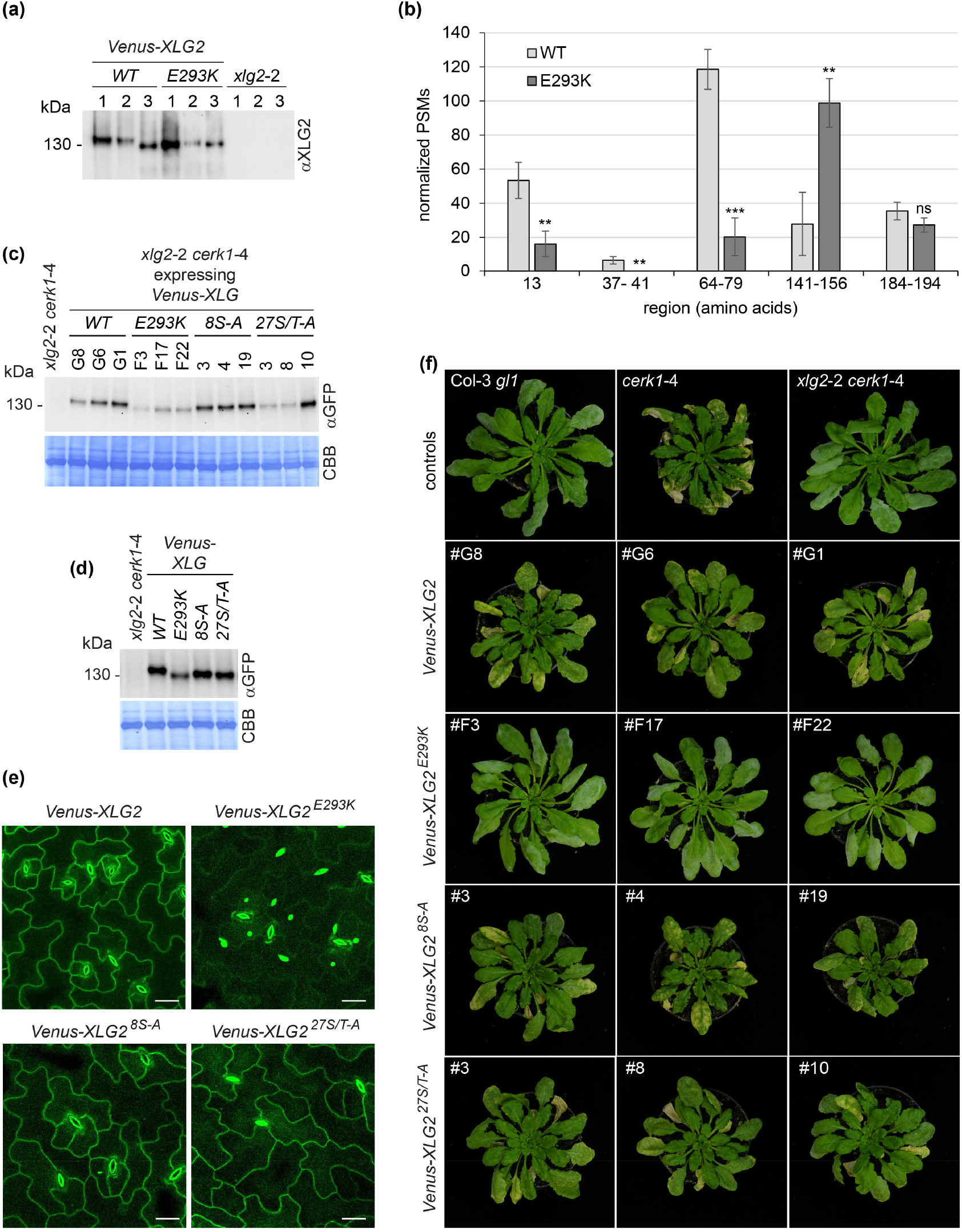
Phosphorylation of the XLG2 N-terminal region affects protein mobility in Western blots, but not XLG2 subcellular localisation and function in the *cerk1*-4 signalling pathway. (a) De-phosphorylation assay: Venus-XLG2 (WT) and Venus-XLG2^E293K^ were purified from *Arabidopsis* plants via the Venus tag and either frozen directly (1), incubated with assay buffer (2) or incubated with assay buffer containing Lambda phosphatase (3). Western blotting was performed with a XLG2-specific antibody and *xlg2*-2 was included as a negative control. (b) Phosphoproteomic analysis of Venus-XLG2 (WT) and Venus-XLG2^E293K^ expressed in the *xlg2*-2 background. Peptide spectrum matches (PSMs) corresponding to phosphorylated peptides were acquired in tSIM mode and normalized to overall XLG2 PSM counts. Numbers of phospho-PSMs covering the indicated amino acid (aa) regions are shown. Data are averages of three independent transgenic lines ±SD. (c-f) *xlg2*-2 *cerk1*-4 plants were transformed with *pXLG2-Venus-XLG2* (WT), *Venus-XLG2*^*E293K*^, *Venus-XLG2*^*8S-A*^ or *Venus-XLG2*^*27S/T-A*^. (c) Western blots of three transgenic lines each, using an anti-GFP/Venus antibody. The genetic background *xlg2*-2 *cerk1*-4 is shown as a negative control. (d) Western blot for direct comparison of SDS-PAGE motility of the indicated fusion proteins. (e) CLSM images of epidermal cells of the indicated transgenic lines. Pictures are maximum projections of 12 focal planes recorded 1μm apart. Size bar = 25 μm. (f) Transgenic lines and controls were inoculated with *E. cruciferarum* and images were taken 14 dpi. Three independent lines are shown per construct.

### XLG2 functions in *cerk1*-4 cell death signalling together with the E3 ubiquitin ligase PUB2

All three *Arabidopsis* XLGs were reported to interact with the E3 ligases PLANT U-BOX 2 (PUB2) and PUB4. Moreover, *xlg1 xlg2 xlg3* triple and *pub2 pub4* double mutants share similar developmental and hormone response defects (Wang *et al*., 2017). Indeed, in our phosphoproteomics analyses, PUB4 and PUB2 were robustly detected as interactors of Venus-XLG2 (WT) as well as Venus-XLG2^E293K^ (Fig. 8a). Immunoblot analysis of *pub4* and *pub2* mutants with a XLG2-specific antibody did not reveal any obvious size differences of XLG2, suggesting that PUB4 and PUB2 do not mediate modification of XLG2 (Fig. 8b). To investigate if PUB4 and PUB2 play a role in the formation of the *cerk1*-4 cell death phenotype, the respective mutants were crossed with *cerk1*-4 and inoculated with *Bgh*. These experiments showed clearly reduced cell death formation in *pub2*-1 *cerk1*-4 compared to *cerk1*-4 alone (Fig. 8c). Similarly, *Bgh*-induced PR1 accumulation was lower in *pub2*-1 *cerk1*-4 than in *cerk1*-4 (Fig. 8d). The *pub4*-3 mutation however was not able to suppress the *cerk1*-4 phenotype (Fig. 8c,d), suggesting that PUB2 is the predominant interactor in this process.

**Figure 8:**
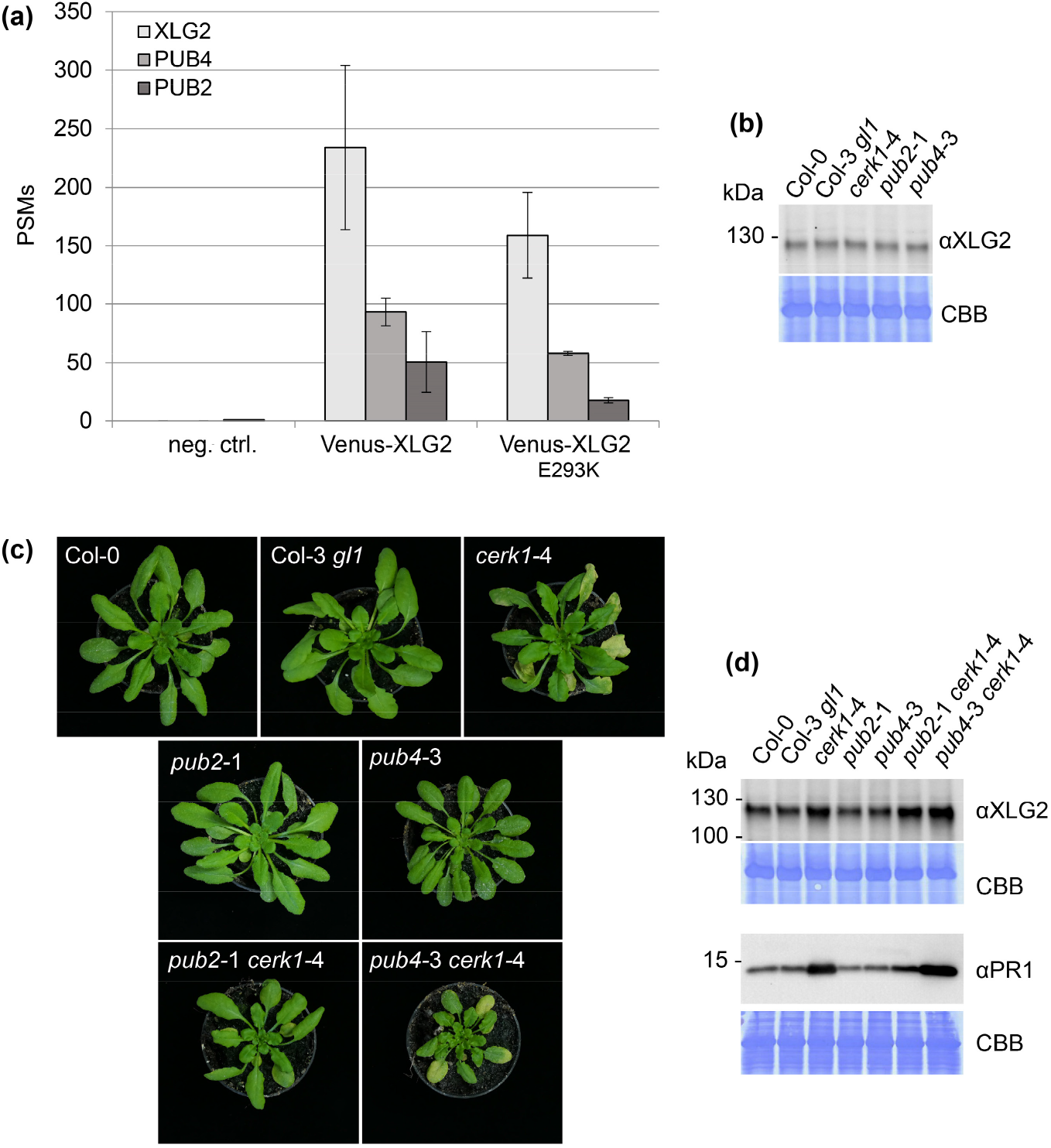
XLG2 functions in *cerk1*-4 cell death signalling together with PUB2. (a) Venus-XLG2 and Venus-XLG2^E293K^ were isolated from stably transformed *Arabidopsis* lines with anti-GFP nanobody beads. The samples were analysed by LC-MS/MS and PSMs matching XLG2/XLG2^E293K^, PUB4 and PUB2 were counted. Data are presented as average ±SD of three independent transgenic lines. Untransformed *xlg2*-2 served as a negative control. (b) Western blot of the indicated lines with a specific XLG2 antibody. (c,d) *pub2*-1 and *pub4*-3 were crossed with *cerk1*-4. Single and double mutants as well as the background accessions Col-0 and Col-3 *gl1* were inoculated with *Bgh* and analysed 7dpi. (c) Images of representative plants. (d) 6 whole rosettes per genotype were pooled and used for Western blots with XLG2 and PR1-specific antibodies. CBB = Coomassie brilliant blue stained membrane.

## DISCUSSION

In this study, we show that loss of function mutations in the heterotrimeric G-protein α-subunit XLG2 suppress the *cerk1*-4 phenotype. *cerk1*-4 plants harbour a single amino acid exchange in the extracellular domain of CERK1 and display excessive cell death and SA accumulation after infection with powdery mildews. Previous analyses showed that in contrast to chitin signalling, this phenotype is independent of CERK1 kinase activity, but correlates with an altered CERK1 shedding status (Petutschnig *et al*., 2014). This suggests a novel pathway, where CERK1-4 generates a cell death signal, but is not itself the signal transducing kinase. Identification of *xlg2* mutations as suppressors confirms the idea that *cerk1*-4-associated cell death involves the generation of a specific molecular signal and active signal transduction by the plant.

In transient expression systems, XLG2 localises to the nucleus, cytosol and plasma membrane (Chakravorty *et al*., 2015; Maruta *et al*., 2015), while XLG2 expressed from its native promoter is found predominantly at the cell periphery in stable transgenic *Arabidopsis* plants. This implies that nuclear localisation of XLG2 reflects a stress response, triggered for example by *Agrobacterium* infiltration. Indeed, when *Arabidopsis* plants were inoculated with spores of the powdery mildew *E. cruciferarum*, (which elicits the runaway cell death phenotype in *cerk1-*4), attacked cells showed nuclear XLG2 signals along with overall increased XLG2 levels. From the *cerk1-*4 suppressor screen, we isolated a non-functional XLG2 variant with an amino acid exchange (E293K) in the N-terminal domain, which showed almost exclusive nuclear localisation. To investigate the relationship between XLG2 subcellular localisation and function in the *cerk1-*4 pathway, we generated XLG2 versions with altered subcellular localisation by appending additional NES and NLS signals or mutating endogenous NES and NLS sequences (XLG2^nes387^ and XLG2^nls426^, respectively). Experiments with these variants revealed that XLG2 localisation is determined by the N-terminal domain and that XLG2 performs its role in *cerk1-*4-associated cell death signalling at the cell periphery. These results suggest that a CERK1-4-derived signal is perceived by a PM-resident sensor, as outlined further below. The role of XLG2 in the nucleus upon fungal infection is currently unclear. (Heo *et al*., 2012) reported that XLG2 interacts with the DNA binding protein RELATED TO VERNALISATION 1 (RTV1) in the nucleus. RTV1 promotes flowering (Heo *et al*., 2012), but there is no evidence for any involvement in plant immunity to date.

The N-terminal domain of XLG2 contains a highly conserved cysteine-rich region with four invariant CxxC motifs. It is reminiscent of a zinc finger domain and therefore has been speculated to possess DNA binding activity (Lee & Assmann, 1999; Ding *et al*., 2008; Heo *et al*., 2012). If the cysteine rich region specifically mediated nuclear functions of XLG2 such as DNA binding, mutation of this segment should not affect *cerk1-*4 signalling. However, disruption of CxxC motifs resulted in non-functional XLG2 proteins with nuclear localisation. These resembled XLG2^E293K^ and XLG2^nls426^, which carry mutations within the N-terminal domain, but not the cysteine rich region. Taken together the results suggest that the CxxC motifs, as well as other conserved regions, are important for the integrity of the entire N-terminal domain, which is in turn essential for XLG2 function.

XLG2 has weak GTP binding / GTPase activity *in vitro* (Heo *et al*., 2012; Lou *et al*., 2020) but it is currently not entirely clear whether this is relevant *in planta*. (Liang *et al*., 2018) reported that XLG2 variants with deleted guanine nucleotide-binding motifs display stabilized interaction with AGB1. Analysis of *xlg2 xlg3* plants expressing the XLG2 deletion variants suggested that GTP binding contributed to a full flagellin-induced ROS burst and resistance to virulent *Pseudomonas* bacteria (Liang *et al*., 2018). On the other hand, a recent study showed that an XLG2 variant which is unable to bind GTP (XLG2^T476N^) had normal interaction with Gβγ and was able to complement PAMP-induced ROS generation and resistance to *Pseudomonas* in *xlg2 xlg3* plants (Maruta *et al*., 2021). Our data indicate that the nucleotide-depleted XLG2^T476N^ (Heo *et al*., 2012; Maruta *et al*., 2021) and the potentially GTPase-dead variant XLG2^R673L^ are functional in *cerk1*-4 signalling. These findings support the idea that XLG2 can fulfil at least some of its functions independently of nucleotide exchange.

XLG2 physically interacts with AGB1 and AGG1/AGG2 at the plasma membrane (Chakravorty *et al*., 2015; Maruta *et al*., 2015) and the resulting heterotrimeric complex signals not only downstream of the flagellin receptor (Liang *et al*., 2016; Liang *et al*., 2018; Wang *et al*., 2018), but also mediates constitutive immune responses in the *bir1* mutant (Liu *et al*., 2013; Maruta *et al*., 2015). This led us to test if AGB1 and AGG1 contribute to powdery mildew-induced excessive cell death in *cerk1-*4. We found that *cerk1*-4 *agb1*-2 and *cerk1*-4 *agg1*-1 had a partially suppressed cell death phenotype with weaker, but not abolished macroscopic lesions. In the case of *cerk1*-4 *agg1*-1, this could be due to an intact AGG2 protein, which we could not address in *cerk1*-4 *agg1*-1 *agg2*-1 mutants because of genetic linkage between *AGG2* and *CERK1*. In contrast, AGB1 does not have any functionally redundant homologs in *Arabidopsis*. Consequently, the results indicate that AGB1 enhances *cerk1-*4 cell death signalling, but is not essential for the process. An explanation for this outcome could be that AGB1 itself does not relay any signals, but affects the status of XLG2. Indeed, Western blot analyses suggested altered phosphorylation of XLG2 in the *agb1*-2 and *agg1 agg2* background. However, since several different multiple phosphorylation mutants of XLG2 still function normally in *cerk1-*4 signalling, it seems most likely that the changed subcellular localisation of XLG2 in *agb1*-2 is the reason for the partial cell death suppression. XLG2 is shifted towards the nucleus in *agb1*-2 plants, suggesting that interaction with the Gβγ dimer helps tether XLG2 to the plasma membrane. As overexpression of Gβγ can sequester XLG3 at the plasma membrane (Chakravorty *et al*., 2015), this seems to be a general mechanism regulating XLG localisation. If Gβγ is absent, less XLG2 might be available for signalling from membrane-associated receptors.

Knockout or mutation of immunity genes can lead to constitutive defense responses and one proposed explanation is guarding by nucleotide-binding leucine-rich repeat (NLR) proteins (Roux *et al*., 2011; Schwessinger *et al*., 2011). This hypothesis has recently gained new momentum by the discovery that *bak1*-4-associated autoimmunity is dependent on helper NLRs (Wu *et al*., 2020). However, the *cerk1-*4 cell death phenotype is conferred by the N-terminal part of the CERK1-4 protein and the kinase domain is not required (Petutschnig *et al*., 2014). This makes an intracellular perception system an unlikely scenario. It is conceivable that CERK1-4 generates a cell death signal in the apoplast, which could be sensed by a RLK or RLP-type receptor that interacts with the XLG2-AGB1-AGG1/AGG2 heterotrimeric G-protein complex. Many receptors characterized to date use BAK1 as a co-receptor and/or SOBIR1 as an adaptor kinase (Escocard de Azevedo Manhaes *et al*., 2021). Analysis of *cerk1-*4 double mutants with *bak1* and *sobir1* showed that the *cerk1-*4 signalling pathway does not rely on these two RLKs. Thus, the signal perception machinery triggering excessive immune responses in *cerk1*-4 and *bir1* mutants (Liu *et al*., 2013; Liu *et al*., 2016) must be different. Interestingly, *cerk1-*4 *bak1* and *cerk1-*4 *sobir1* double mutants even had increased *PR1* expression after powdery mildew infection, suggesting that BAK1 and/or SOBIR1-containing immune complexes compete for shared downstream signalling components (such as XLG2) with the receptor system that senses CERK1-4. Competitive binding of different RLKs to XLG2 was recently reported (Maruta *et al*., 2021).

XLG2 has been shown in multiple studies to facilitate PAMP-induced ROS bursts together with XLG3 (Maruta *et al*., 2015; Liang *et al*., 2016; Liang *et al*., 2018; Maruta *et al*., 2021). Therefore, we investigated if the role of XLG2 in ROS control is important for the formation of the *cerk1*-4 phenotype. We found that XLG2 mediates flagellin-triggered ROS production redundantly with not only XLG3, but also XLG1. Since the *cerk1*-4 phenotype is suppressed by *xlg2* alone, this argued against an involvement of ROS in *cerk1*-4 cell death formation. Indeed, XLG2 variants with mutated phosphorylation sites that are compromised in ROS regulation (Liang *et al*., 2016) were fully functional in the *cerk1*-4 pathway. These findings were corroborated by the fact that *rbohD* and *rbohF* mutants were unable to suppress the *cerk1*-4 phenotype.

XLG2 in *agb1*-2, the XLG2^E293K^ variant and all other non-functional XLG2 versions generated in this study showed faster migration in SDS-PAGE than the wild type version of the protein. Phosphatase assays indicated that the difference in apparent mass was caused by phosphorylation, therefore we quantitatively analysed phosphorylation patterns on XLG2 and XLG2^E293K^. Interestingly, sites required for proper ROS induction (Liang *et al*., 2016) were more phosphorylated in XLG2^E293K^ than the wild type. Liang *et al*. (2016) found S141, S148, S150 and S151 to be phosphorylated in flagellin-challenged cells. Therefore, it is worth noting that we found phosphorylation in this region in the absence of any PAMP treatment. Residues closer to the N-terminus (S/T 13-79) were significantly less phosphorylated in XLG2^E293K^ than wild type XLG2, therefore we generated corresponding XLG2 mutation variants. Analysis of *Arabidopsis* lines stably expressing these constructs confirmed that the lower apparent mass of XLG2^E293K^ is caused by reduced phosphorylation, but also showed that this is not the reason why XLG2^E293K^ is non-functional. Presumably, XLG2 is not regulated by phosphorylation in the *cerk1*-4 pathway. XLG2^E293K^ is unable to interact with AGB1, which can partially explain its lack of function, but it seems very likely that the E293K mutation also disrupts interaction with other proteins.

In our mass spectrometry analyses, we robustly co-purified PUB2 and PUB4 together with XLG2. Both ubiquitin E3 ligases were previously reported to interact with XLGs and function in cytokinin signalling and floral development (Wang *et al*., 2017). We isolated PUB2 and PUB4 with wild type XLG2 as well as the E293K variant. This suggests that they interact with XLG2 at the cell periphery and the nucleus and that the interaction may not be dependent on an intact N-terminal domain. Western blot analysis of XLG2 in *pub2* and *pub4* mutants did not show any apparent mass differences to the wild type background. Thus, these E3 ligases might not modify XLG2, but might be regulated by XLG2 to modify other proteins. Mainly PUB4 is required for full chitin induced ROS generation (Desaki *et al*., 2019; Derkacheva *et al*., 2020) and it is tempting to speculate that this function depends on XLG2/3. Interestingly, *pub2* but not *pub4* was able to partially suppress powdery mildew induced cell death in *cerk1*-4. It seems that both PUBs interact with XLGs in many different pathways, but there is some level of specialisation between them.

Taken together, our results show that XLG2 functions in a novel, so far uncharacterized pathway to transduce the CERK1-4-specific cell death signal at the plasma membrane. In contrast to previously described immune signalling systems that utilize XLG2, this mechanism is independent of BAK1 and SOBIR1 and not regulated by NADPH-generated ROS or XLG2 phosphorylation in the N-terminal domain.

## ACKNOWLEDGEMENTS

We thank Jian-Min Zhou (Institute of Genetics and Developmental Biology, Chinese Academy of Sciences) for providing XLG2-FLAG, XLG2 4S-A-FLAG and XLG2 4S-D-FLAG constructs. We especially thank Sabine Wolfarth and Ludmilla Heck-Hrarti for technical assistance, and Kerstin Schmitt and Oliver Valerius (Core Facility LCMS Protein Analytics of the Faculty of Biology and Psychology and the Faculty of Forest Sciences and Forest Ecology, Georg-August University Göttingen) for mass spectrometry analyses. This work was funded by the DFG grants LI 1317/10-1 and INST 186/1277-1 FUGG.

## Supporting Information

### SUPPLEMENTAL MATERIALS AND METHODS

#### Mass Spectrometry

After GFP pull-down, on bead trypsin digestion was performed as described in a protocol provided by the manufacturer Chromotek, which is based on (Hubner *et al*., 2010). Digested peptides were purified with the C18 stop and go extraction (Stage) method (Rappsilber *et al*., 2007). Purified peptides were vacuum dried at 45°C and then dissolved in 20 μl 2% acetonitrile, 0.1% formic acid.

3-4 μl of each sample were subjected to reversed-phase liquid chromatography for peptide separation using an RSLCnano Ultimate 3000 system (Thermo Fisher Scientific). Peptides were loaded on an Acclaim^*®*^ PepMap 100 pre-column (100 μm × 2 cm, C18, 5 μm, 100 Å; Thermo Fisher Scientific) with 0.07 % trifluoroacetic acid. Analytical separation of peptides was done on an Acclaim^*®*^ PepMap RSLC column (75 μm × 50 cm, C18, 2 μm, 100 Å; Thermo Fisher Scientific) running a water-acetonitrile gradient at a flow rate of 300 nl/min. All solvents and acids had Optima grade for LC-MS (Thermo Fisher Scientific). Chromatographically eluting peptides were on-line ionized by nano-electrospray (nESI) using the Nanospray Flex Ion Source (Thermo Fisher Scientific) at 1.5 kV (liquid junction) and continuously transferred into the mass spectrometer (Q Exactive HF, Thermo Fisher Scientific). Full scans in a mass range of 300 to 1650 m/z were recorded at a resolution of 30,000 followed by data-dependent top 10 fragmentation (HCD) at a resolution of 15,000 (dynamic exclusion enabled).

For targeted single-ion monitoring (tSIM) LC-MS runs, the resolution was set to 60,000. The maximum ion time for tSIM scans (AGC target 1e6) was 100 ms. The loop count equalled the number of m/ z values on the inclusion list. Dynamic exclusion was disabled. LC-MS method programming and data acquisition was performed with the XCalibur software 4.0 (Thermo Fisher Scientific).

Database searches were performed using Proteome Discoverer 2.2 software (Thermo Fisher Scientific) with the Mascot and SequestHT search algorithms against the Araport11 protein database (Cheng *et al*., 2017). The digestion mode was set to trypsin and the maximum number of missed cleavage sites was set to three. Carbamidomethylation of cysteines was set as fixed modification, oxidation of methionine and serine, threonine and tyrosine phosphorylation were set as variable modifications. The mass tolerance was 10 ppm for precursor ions and 0.02 Da for fragment ions. The decoy mode was rev with a false discovery rate of 0.01.

Phosphopeptides were identified in initial runs and then selected for quantitative analysis by tSIM. Phosphorylation sites within phosphopeptides were assigned with the ptmRS node of Proteome Discoverer 2.2 (Thermo Fisher Scientific). Many partially overlapping phosphopeptides were found and unambiguous identification of phosphorylation sites could be achieved for approximately 50% of obtained spectra. Therefore, phosphopeptides were grouped into distinct, non-overlapping clusters. Phosphorylation within these was quantified by counting phosphorylated peptide spectrum matches (PSMs) and normalizing them to all PSMs obtained for XLG2.

#### Cloning

For generation of complementation constructs, a genomic fragment containing the *XLG2* coding region and 1260 bp of upstream sequence was amplified with primers EP209 and EP210 (Table S1). The PCR product was cloned into a pGreenII-0229 (Hellens *et al*., 2000) derivative that contains a modified MCS and the 35S terminator, via its *Asc*I and *Bam*HI sites. pGreenII-0229*-pXLG2-Venus-XLG2 and XLG2*^*E293K*^ were cloned by homologous recombination in yeast. PCR products of *pXLG2, Venus* and the *XLG2/XLG2*^*E293K*^ coding region were amplified with primer pairs CM82 and CM92, JE23 and CM93, CM94 and CM83 respectively (Table S1). The primers introduced overlaps between the PCR products and with the shuttle vector pRS426 (Christianson *et al*., 1992) as well as *Asc*I and *Bam*HI sites for downstream cloning. *Saccharomyces cerevisiae* strain S288c BY4741 (Brachmann *et al*., 1998) was transformed with a mix of PCR fragments to be recombined and pRS426 linearized with *Bam*HI and *Kpn*I. Cells were then plated on SC plates (-Ura +Gluc) and incubated at 28°C for 2 – 3 days. Transformed *S. cerevisiae* cells were washed from plates with water, pelleted by centrifugation (5min, 4000g, RT), resuspended in 200μl P1 buffer of the QIAGEN Plasmid Midi kit and disrupted by incubation on a vibrax with 0.3g glass beads (425-600 micron) for 10 - 15 minutes. Subsequently, glass beads were pelleted by centrifugation and plasmids were prepared according to the kit manual. The pRS426 vector construct was transformed into *E. coli* TOP10 cells for amplification. Next, the plasmid was isolated and cut with *Asc*I and *Bam*HI for cloning into a modified pGreenII-0229 vector. The mutations nls426, nes387, C214/217A, C229/232/237/240A and C254/257A were introduced to pGreenII-0229-*pXLG2-Venus-XLG2* using the NEB Q5m Site Directed Mutagenesis kit and primers listed in Table S1. In order to add c-terminal NLS/NES/nls/nes sequences, an additional *Not*I site was introduced after the *XLG2* coding region by site directed mutagenesis with primers JF17 and JF18. The resulting vector was linearized with *Not*I and *Bam*HI and ligated with annealed oligonucleotide pairs JF19 and JF20, JF21 and JF22, JF30 and JF31 or JF32 and JF33 (Table S1). For the 8S-A and 27S/T-A mutations, synthetic DNA fragments containing the mutations were purchased (Thermo Fisher Scientific) and amplified with primers EP624 and EP629 (Table S1). A PCR on pGreenII-0229-*pXLG2-Venus-XLG2* was carried out with primers EP625 and EP220 (Table S1) and the two fragments were joined with Gibson assembly.

All constructs were confirmed by sequencing, transformed into *Agrobacterium tumefaciens* GV3101 (pSoup) and used for stable transformation of *Arabidopsis* plants.

## SUPPLEMENTAL FIGURES and FIGURE LEGEND

**Figure S1:**
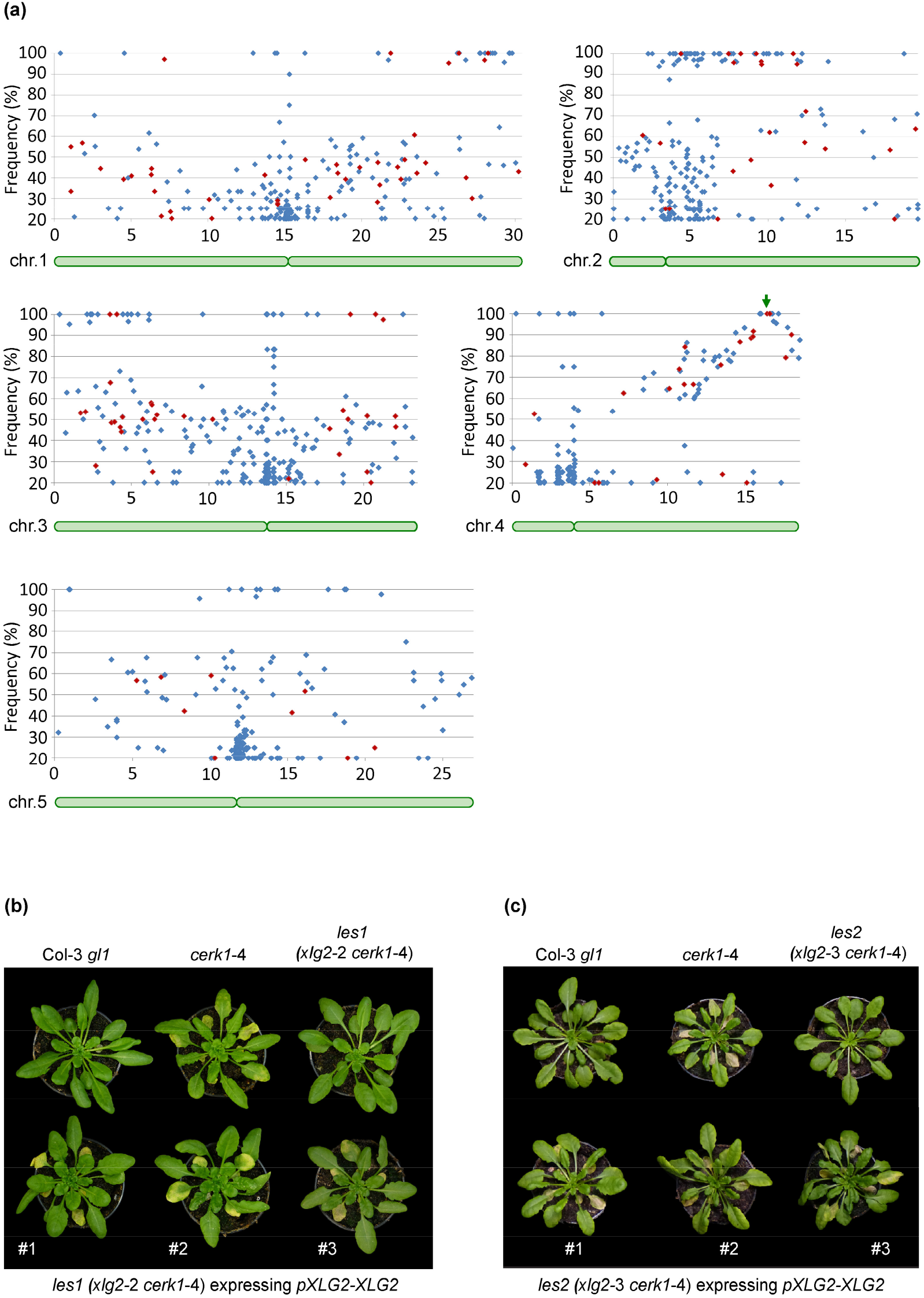
Mapping and complementation of *les* mutations. (**a**) *les1* was backcrossed to *cerk1-*4 and in the F2 generation, powdery mildew-induced macroscopic cell death was scored. DNA of 31 suppressor plants was pooled and Illumina sequenced. Reads were aligned to the TAIR10 genome and SNPs identified. The frequencies of *les1*-specific SNPs were plotted against their position on the respective chromosomes (x-axis unit = 10^6^ bp). Synonymous mutations are shown as blue and non-synonymous mutations as red diamonds. The causative *les1* mutation in *At4g34390* is marked with a green arrow and the chromosome arms are represented by green bars. **(b, c)** *les1 (xlg2-*2 *cerk1-*4) and *les2 (xlg2-3 cerk1-*4) plants were transformed with a genomic DNA fragment containing the wild type *XLG2* gene including its promoter. The resulting transgenic plants were inoculated with *Bgh* and images were taken after 7 or 10 days. The pictures show three independent transgenic lines as well as Col-3 *gl1, cerk1-*4 and *les1* or *les2* control plants. (b) and (c) are composite figures of 6 individual images.

**Figure S2:**
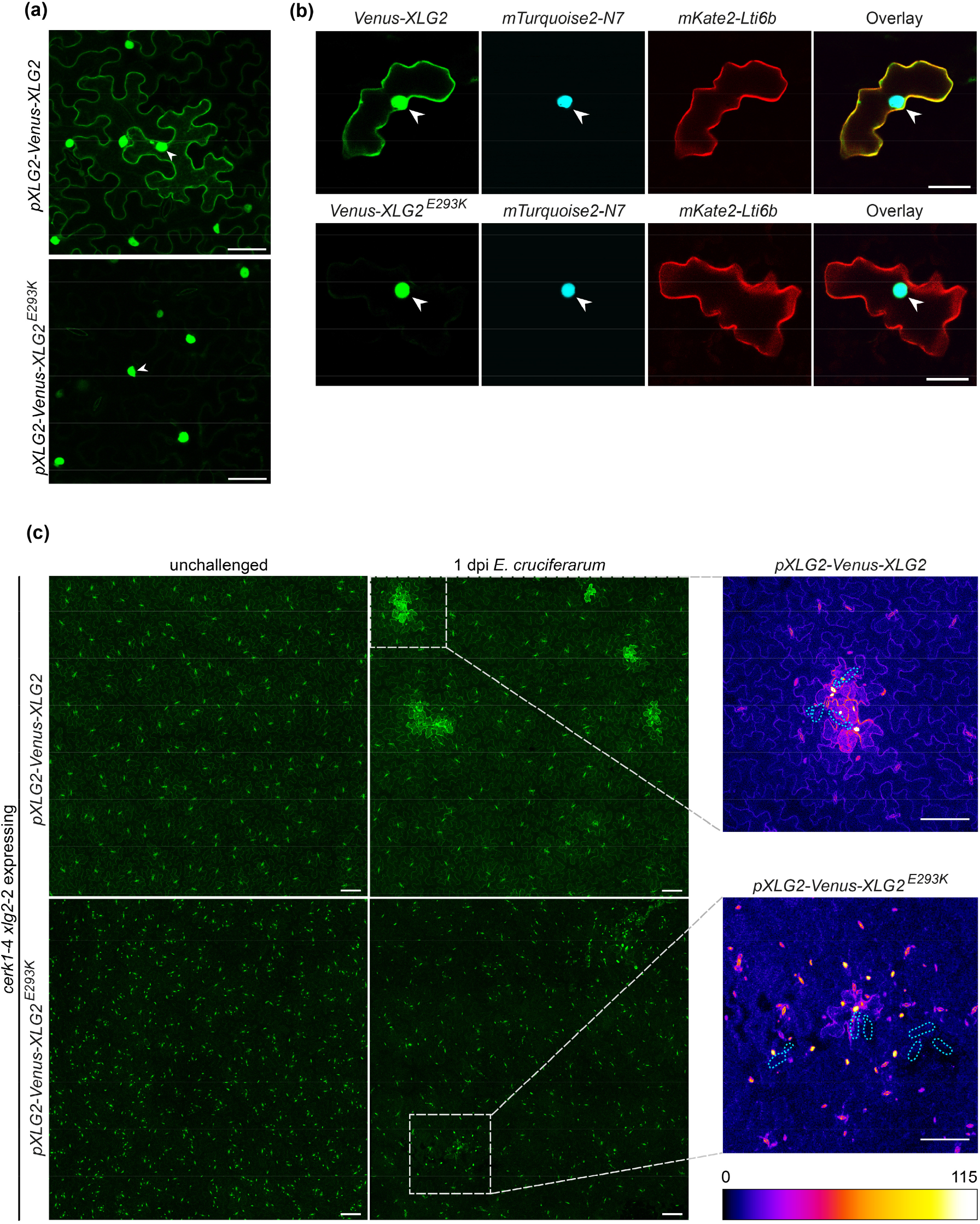
Localisation of Venus-XLG2 and Venus-XLG2^E293K^ in transient expression systems and stably transformed *Arabidopsis* plants. (a) Venus-XLG2 WT and Venus-XLG2^E293K^ were transiently expressed in *N. benthamiana* and CLSM was performed 3dpi. Representative images are shown as maximum projections of 31 focal planes recorded 1μm apart. Scale bar = 50μm. (b) Particle bombardment in *Arabidopsis xlg2*-2 plants. Venus-XLG2 and Venus-XLG2^E293K^ were co-transformed with Turquoise2-N7 (nuclear marker) and mKate2-LTI6b (PM marker). Epidermal cells were analysed 2d after bombardment and representative single plane images are shown. Scale bar = 50μm. Nuclei are highlighted with arrowheads. (c) Leaf areas of transgenic *Arabidopsis* plants expressing Venus-XLG2 or Venus-XLG2^E293K^ in the *cerk1*-4 *xlg2*-2 background, unchallenged and 1dpi with *E. cruciferarum*. Images are stitched maximum projections of 60 focal planes recorded 1.5 μm apart. Scale bar = 100 μm. On the right, zoom-ins of areas marked with dashed boxes are shown. Fungal spores are outlined with dashed blue lines. For better visualization of fluorescence levels, a multicolor lookup table was applied. Scale bar = 100 μm.

**Figure S3:**
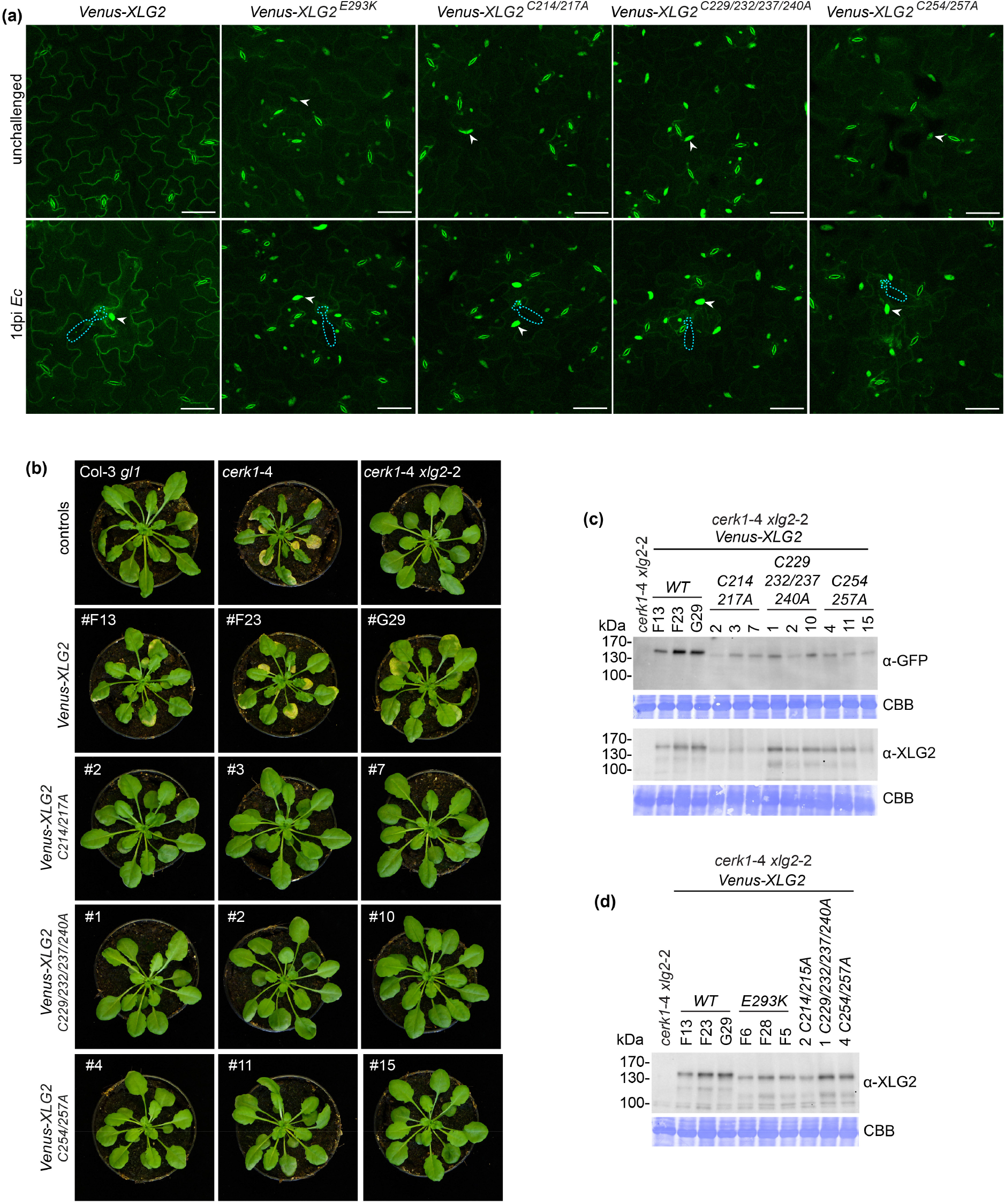
Signalling of XLG2 in the *cerk1*-4 pathway requires an intact cysteine-rich region. Stably transformed *cerk1*-4 *xlg2*-2 lines expressing *Venus-XLG2, Venus-XLG2*^*E293K*^ or *Venus-XLG2* with the indicated C to A mutations, were analysed. (a) Representative CLSM images of unchallenged and *E. cruciferarum*-infected plants (1 dpi). Pictures are maximum projections of z-stacks spanning the epidermal cell layer. Example nuclei are marked by arrowheads and the positions of fungal spores at sites of attempted penetration are outlined by dashed blue lines. Scale bar = 50 μm. (b) *cerk1*-4 *xlg2*-2 lines expressing the same constructs and non-transformed controls were inoculated with *E. cruciferarum* and images were taken 14 dpi. Three representative lines are shown. (c) Accumulation of Venus-XLG2 was confirmed by Western blotting after *E. cruciferarum* infection. (d) Apparent molecular masses of Venus-XLG2 wild type, XLG2^E293K^ and the C to A mutant versions were compared in Western blots by running them side by side. CBB = Coomassie brilliant blue stained membrane.

**Figure S4:**
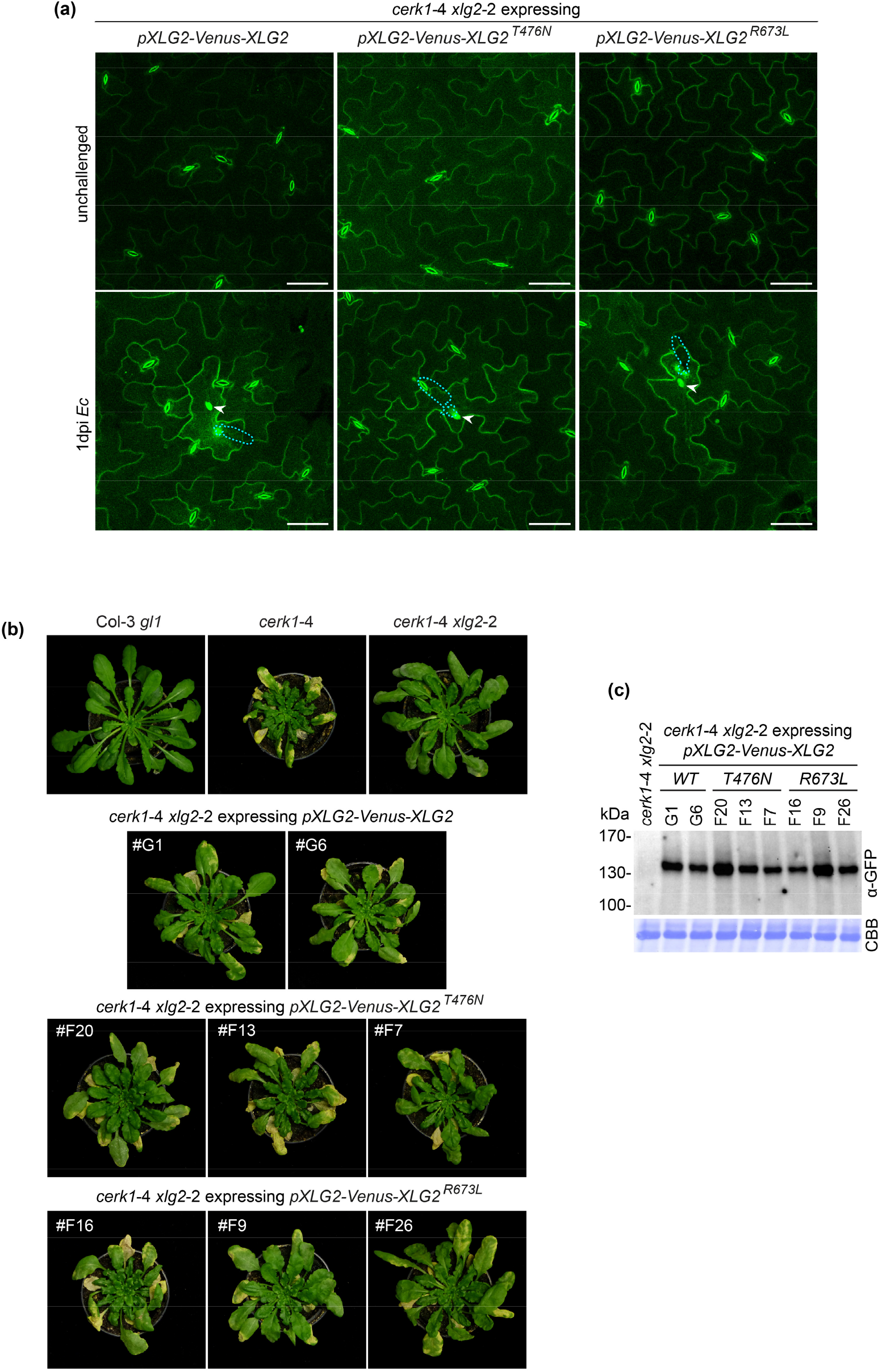
XLG2 GTP binding and GTPase activity are not required for *cerk1-*4 phenotype formation. Transgenic *cerk1*-4 *xlg2*-2 lines *expressing pXLG2-Venus-XLG2, Venus-XLG2*^*T476N*^ or *Venus-XLG2*^*R673L*^ were characterized (a) Representative CLSM images of unchallenged and infected (1 dpi *E. cruciferarum*) plants. Pictures are maximum projections of z-stacks spanning the epidermal cell layer. Nuclei in attacked cells are marked by arrowheads and the positions of fungal spores are outlined by dashed blue lines. Scale bar = 50 μm. (b) The indicated transgenic lines and controls were inoculated with *E. cruciferarum* and the macroscopic phenotype was documented 14dpi. Images of two or three independent lines are shown. (c) Western Blots detecting Venus-XLG2 in the lines presented in (b) after *E. cruciferarum* infection. CBB = Coomassie Brilliant Blue stained membrane.

## SUPPLEMENTAL TABLES

**Table S1:**
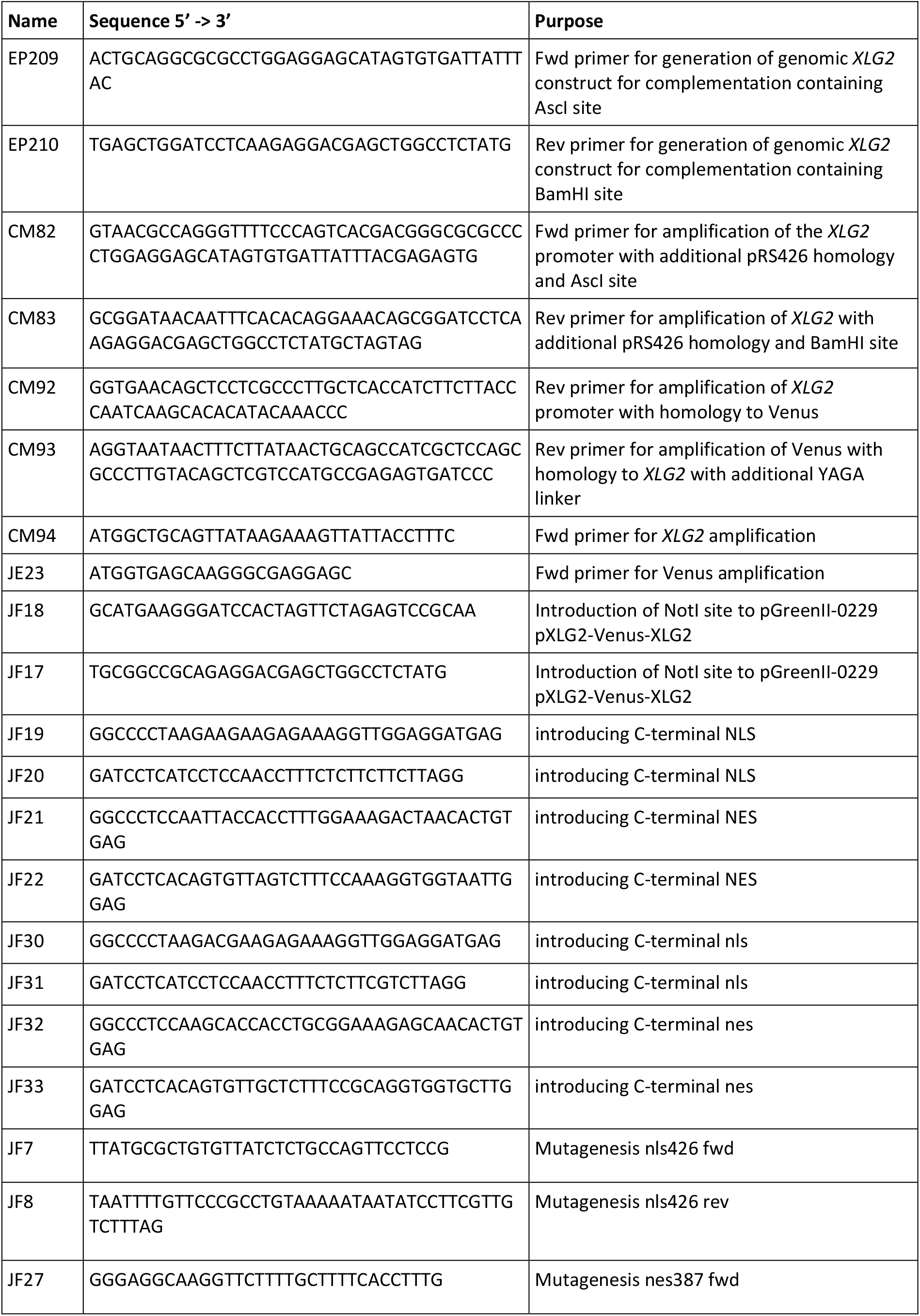
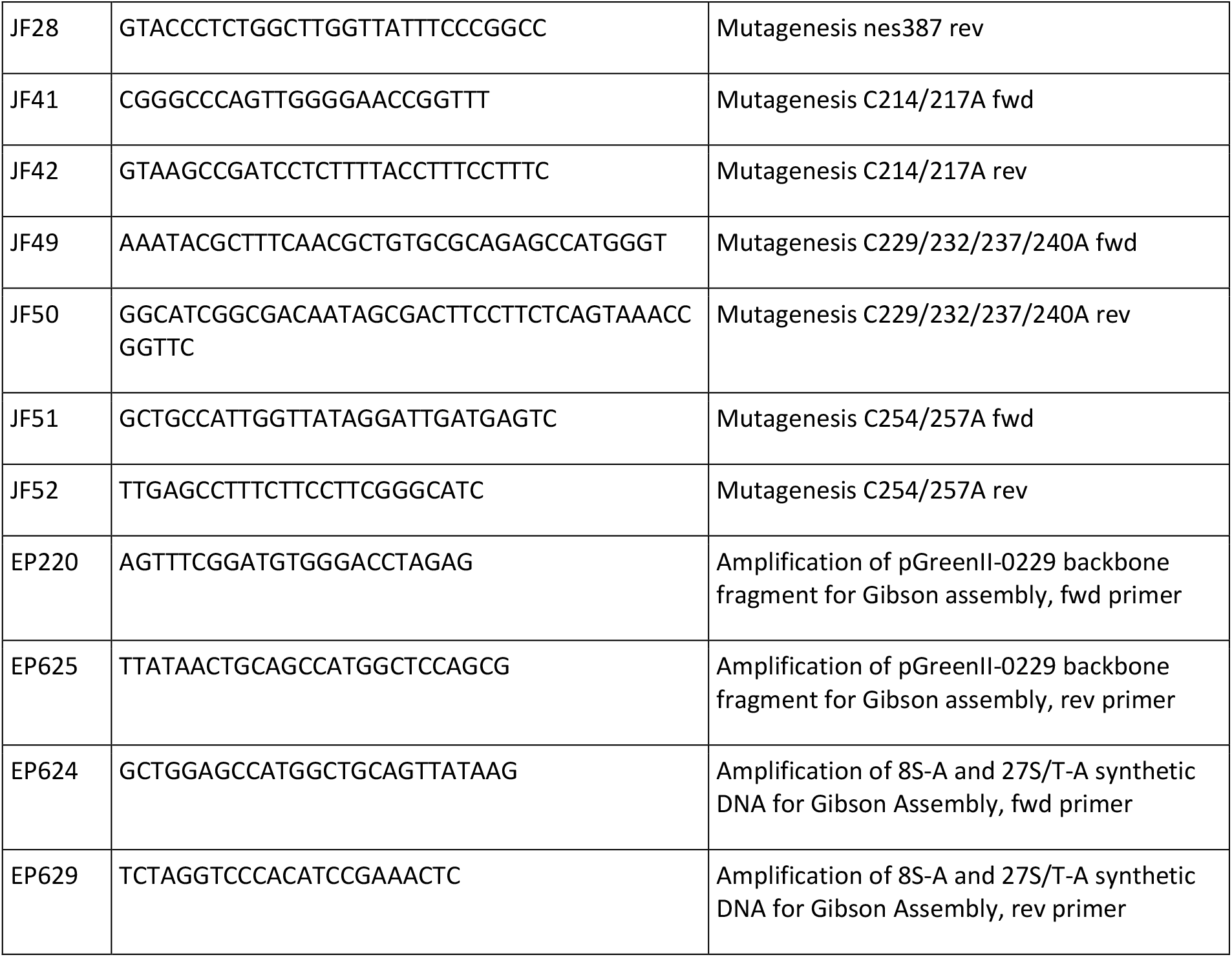
Oligonucleotides used in this study.

**Table S2:** Overview of transgenic XLG2-FLAG lines in the *cerk1*-4 *xlg2-*2 background. ^*^Expression of fusion proteins confirmed via Western blotting. Phenotype evaluation after *E. cruciferarum (Ec)* infection in T1 generation.

| Transgene<br>(expressed from 35S promoter) | Independent lines<br>generated | Independent lines<br>analysed*<br>(complementation rate) | Apparent mass in<br>Western Blots |
| --- | --- | --- | --- |
| *XLG2-FLAG WT* | 18 | 18 (100%) | ~110kDa |
| *XLG2<sup>4S-A</sup>-FLAG* | 20 | 20 (100%) | WT-like |
| *XLG2<sup>4S-D</sup>-FLAG* | 16 | 16 (100%) | WT-like |

**Table S3:** Overview of transgenic Venus-XLG2 lines in the *xlg2-*2 background. ^*^Expression levels and localisation of Venus fusion proteins confirmed via CLSM and/or Western blotting. Phenotype evaluation after *E. cruciferarum (Ec)* infection in T1 and/or T2 generation. All lines showed a WT-like growth and cell death phenotype. ** In lines with very high expression levels, faint nuclear signals were observed.

| Transgene<br>(expressed from<br>XLG2 promoter) | Independent<br>lines<br>generated | Independent<br>lines<br>analysed* | Subcellular<br>localisation in<br>unchallenged<br>cells | Subcellular<br>localisation *Ec*<br>infected cells | Apparent<br>mass in<br>Western<br>Blots |
| --- | --- | --- | --- | --- | --- |
| *Venus-XLG2 WT* | 42 | 6 | cell periphery** | nucleus, cell<br>periphery,<br>cytoplasmic strands | ~140kDa |
| *Venus-XLG2<sup>E293K</sup>* | 15 | 6 | nucleus, very<br>faint cell<br>periphery | nucleus, very faint<br>cell periphery | reduced |

**Table S4:**
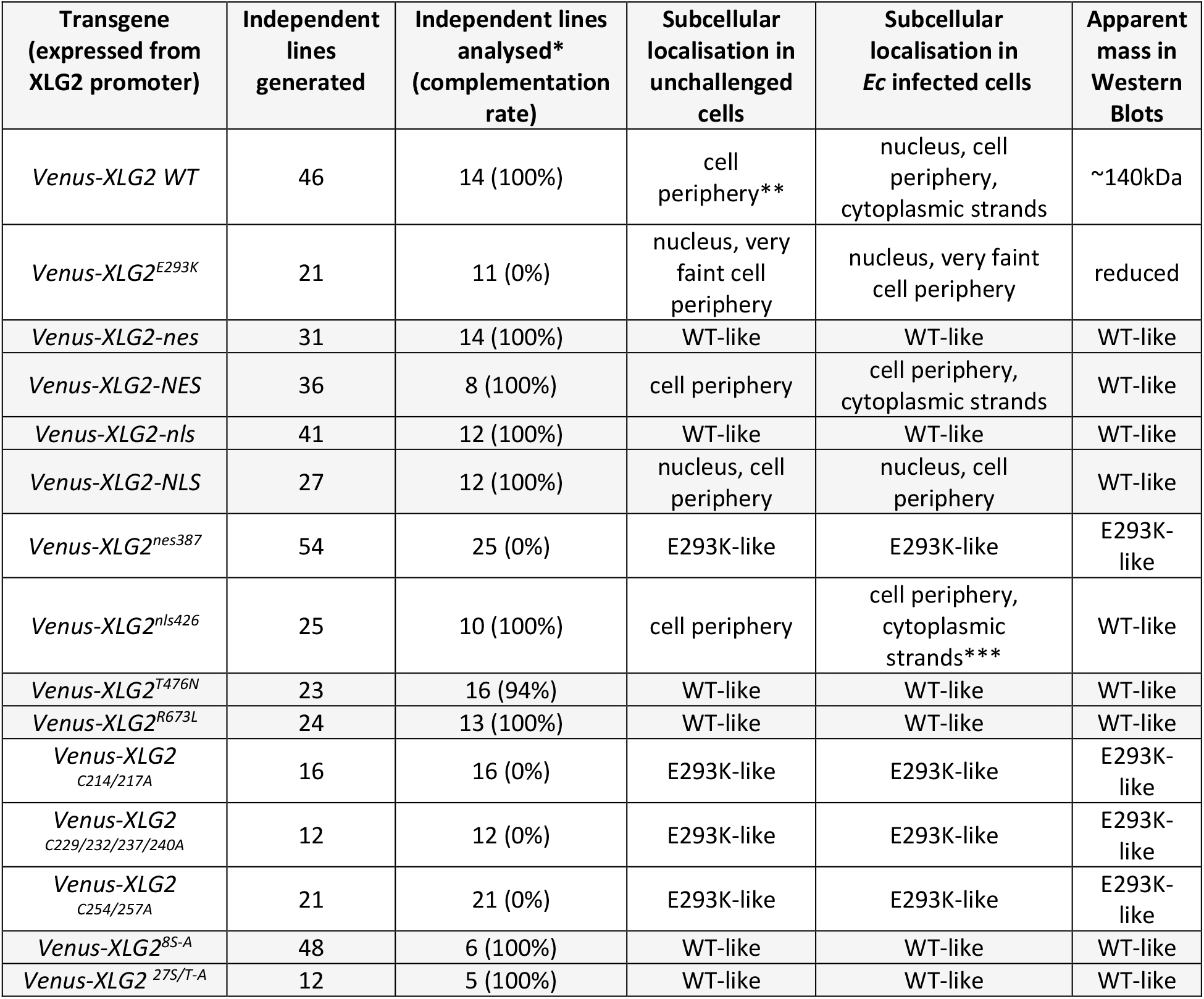
Overview of transgenic Venus-XLG2 lines in the *cerk1*-4 *xlg2-*2 background. Expression levels and localisation of Venus fusion proteins confirmed via CLSM and/or Western blotting, Phenotype evaluation after *E. cruciferarum (Ec)* infection in T1 and/or T2 generation. ** In lines with very high expression levels, faint nuclear signals were observed. *** Occasionally very faint nuclear signals were observed in *E. cruciferarum*-attacked cells.

**Table S5:**
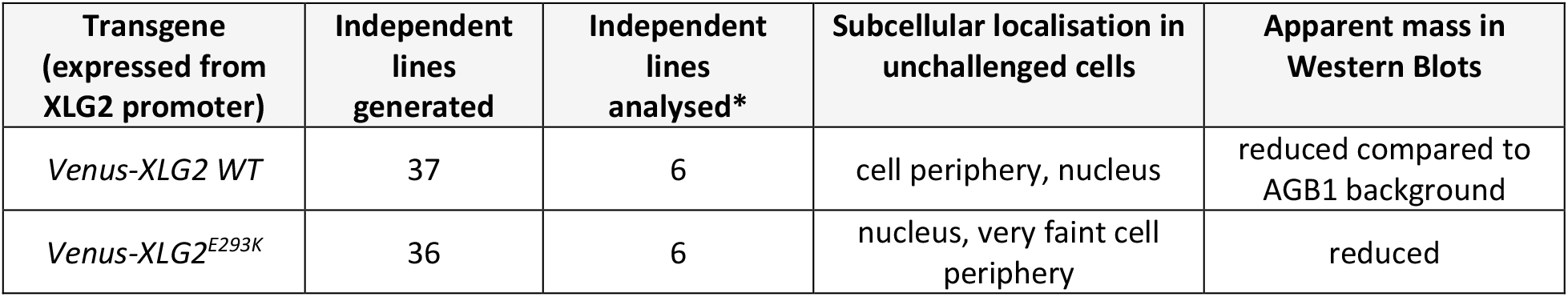
Overview of transgenic Venus-XLG2 lines in the *agb1-2* background. ^*^Expression levels and localisation of Venus fusion proteins confirmed via CLSM and Western blotting.

| Transgene<br>(expressed from<br>XLG2 promoter) | Independent<br>lines<br>generated | Independent<br>lines<br>analysed* | Subcellular localisation in<br>unchallenged cells | Apparent mass in<br>Western Blots |
| --- | --- | --- | --- | --- |
| *Venus-XLG2 WT* | 37 | 6 | cell periphery, nucleus | reduced compared to<br>AGB1 background |
| *Venus-XLG2<sup>E293K</sup>* | 36 | 6 | nucleus, very faint cell<br>periphery | reduced |

## Notes

### Competing Interest Statement

The authors have declared no competing interest.

